# *In vivo* mRNA delivery to the lung vascular endothelium by dicationic Charge-Altering Releasable Transporters

**DOI:** 10.1101/2025.04.19.649649

**Authors:** Mahmoud M. AbdElwakil, Jeffery Ni, Summer Ramsay-Burrough, Paul Joshua Hurst, Rebecca L. McClellan, Samuel R. Khasnavis, Yuan Jia, Sehar R. Masud, Timothy R. Blake, Adrienne Sallets, Ole A. W. Haabeth, Idit Sagiv-Barfi, Debra K. Czerwinski, Ronald Levy, Paul A. Wender, Maya E. Kumar, Robert M. Waymouth

**Affiliations:** Department of Chemistry, Stanford University, Stanford, CA 94305, USA; Division of Oncology, Department of Medicine, Stanford Cancer Institute, Stanford University, Stanford, CA 94305, USA; Department of Chemical and Systems Biology, Stanford University, Stanford, California 94305, USA; Department of Pediatrics, Division of Pulmonary Medicine, Stanford University School of Medicine, USA

**Keywords:** targeted mRNA delivery, degradable polymers, lung endothelium, CryoEM nanoparticles

## Abstract

Endothelial cells (EC) comprise the pulmonary vascular bed and play a significant role in health and disease. Consequently, the EC niche represents an attractive therapeutic target for treating a wide range of pulmonary vascular diseases. We have identified a new class of dicationic Charge-Altering Releasable Transporters. These single-component transporters selectively deliver mRNA to the lung upon intravenous administration without the use of a targeting ligand. Significantly, the number and spatial array of cationic charges within the repeating units of the CART polymer are found to control both mRNA delivery efficacy and tissue tropism. High-resolution imaging revealed efficient mRNA delivery to endothelial cells in pulmonary arteries, veins and capillaries. The selective lung tropism of these new CARTs, coupled with the efficient and tunable synthesis of this new family of CART amphiphiles, represents an enabling platform for research and clinical applications.

## Introduction

The emergence of RNA-based therapies, marked by the widespread adoption of SARS-CoV-2 vaccines created by Moderna and BioNTech/Pfizer, is transforming the landscape of disease prevention, management, and eradication.^1–5^ A key to realizing the full potential of this technology is the selective and efficient delivery of mRNA to the cell types and organs relevant to the disease of interest. However, the control of organ and cell selectivity of nanoparticle delivery is a complex problem^6–10^ involving physicochemical properties like size,^11–12^ shape,^12–13^ surface charge,^14–15^ and surface functionalization^7,16–18^ (e.g. targeting ligands and post-administration protein corona effects). Among the efforts to develop cell– and tissue-specific targeting strategies,^19–25^ delivery vectors that target lung endothelial cells^26–41^ are of particular importance. The pulmonary endothelium is a vital interface between the circulation and alveolar air spaces, facilitating the transfer of gases, liquids, and cells between the blood and the airways. This diverse and highly specialized cellular layer is instrumental in preserving healthy respiratory physiology, regulating inflammatory processes, and coordinating pulmonary immune responses. Impairment of endothelial function within the lungs may lead to the development of several respiratory pathologies, such as acute respiratory distress syndrome, pulmonary arterial hypertension, and complications associated with COVID-19 infection. Because of the intimate association between the pulmonary endothelium and respiratory epithelium, lung-selective nucleic acid delivery would enable new treatments for wide range of lung diseases including infections caused by airborne pathogens,^42–43^ chronic inflammatory diseases,^44–45^ pulmonary fibrosis^46^ and lung cancer,^47^ many of which are poorly addressed or have no cure.

Herein we describe a new class of Charge-Altering Releasable Transporters (CARTs) derived from dicationic amino acids that exhibits exquisite lung-selective mRNA delivery (Figure 1A). CARTs are single-component delivery vectors that have been shown to deliver a variety of nucleic acid cargoes (mRNA, siRNA, pDNA, circRNA) *in vitro* and *in vivo.*^48–58^ CARTs are amphipathic diblock oligomers consisting of an oligocarbonate lipid block followed by a degradable cationic polyester block derived from α-amino acids (e.g. N-hydroxyethylglycine,^48,51^ N-hydroxyethyllysine,^57^ or serine^52^). CARTs are designed to have dynamic electrostatic properties (charge state and oligomeric composition) that change at tunable rates via irreversible degradation of the cationic polyester block through hydrolysis or rearrangement, thereby producing small molecule byproducts and enabling RNA release.^48,57,59^ This charge-altering behavior depends on both pH (CARTs are stable in water at low pH (pH≈5) but degrade at higher pH (pH 6-8)) and the structure of the cationic block.^48,57,50^ The lipid block is a key component of CART polymers; variations in the nature of the side-chain lipid results in CARTs with enhanced *in vitro* transfection of lymphocytes and improved *in vivo* spleen selective mRNA delivery.^23,51^ Moreover, the nature of the cationic block, essential for complexation of anionic nucleic acids, has potential to elicit new *in vivo* tropisms,^60^ as the tissue selectivity upon IV administration is known to be influenced by the nature of the cationic block^61^. Lysine-derived CART amphiphiles, bearing a pendent amino group and two cationic amines per repeat unit, were shown to exhibit high selectivity for protein expression in the lung upon IV administration.^57^ As charge density is known to be important in coacervate phase separation,^62^ here we investigate the role of subtle changes in the nature of the pendent side chain and spacing between CART backbone and pendent amino group with a series of CART amphiphiles derived from the N-hydroxyethyl α-amino acids lysine, ornithine, diaminobutyric acid (Daba) and diaminopropionic acid (Dapa).^63–64^ A series of dicationic CARTs which feature pendent amino groups with a series of carbon spacers between the amine and backbone were made and evaluated for *in vitro* and *in vivo* RNA delivery. First, we synthesized ornithine-derived CARTs (Orn-CARTs) with an amino side chain only one carbon shorter than that of lysine. Transfection assays showed efficient cellular uptake and mRNA translation. Interestingly, in vivo mRNA delivery with Orn-CARTs led to significantly higher levels of protein expression in the lung when compared to mRNA delivery with lysine-derived Lys-CARTs or commercially available polyethyleneimine reagents (*in vivo*-jetPEI^®^). We further expanded the design space of dicationic monomers by incorporating a diaminobutyric acid derived monomer (Daba) with two carbons in the pendent amino group and diaminopropionic acid derived monomer (Dapa) with one carbon in the pendent amino group. Daba-CART and Dapa-CART led to significantly higher levels of protein expression than Orn-CART, but only Daba-CART retains lung selectivity. In-depth characterization of Orn-CART and Daba-CART using transgenic mice and confocal microscopy showed that this family of dicationic CARTs selectively and efficiently deliver mRNA into endothelial cells of the pulmonary arteries, veins and capillaries as well as the lung’s systemic bronchial vessels. This study shows that the spatial array of dicationic subunits in the CARTs have a significant influence on the *in vivo* mRNA delivery efficiency and tissue tropism with potential to advance basic research and address unmet medical needs in the lung vascular endothelium.

**Figure 1.**
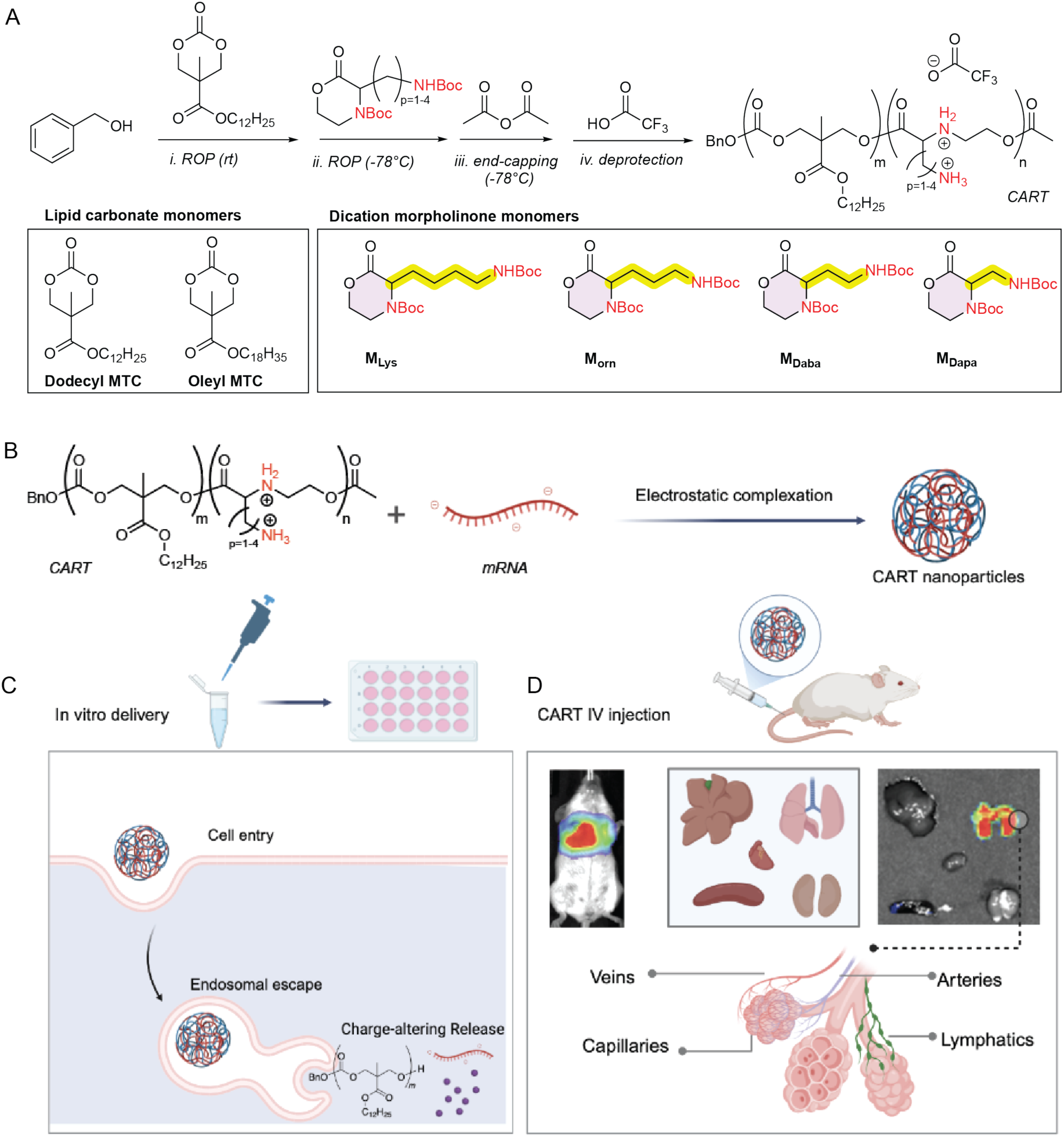
(A) Synthesis scheme and chemical structures of dicationic CARTs: *(i)* room temperature ring-opening polymerization (ROP) of dodecyl MTC using TBD as a catalyst, followed by *(ii)* low-temperature ROP of morpholinone monomer, *(iii)* end-capping with acetic anhydride, and *(iv)* Boc deprotection with TFA. (B) CART polymers complex anionic oligonucleotides (such as mRNA) to form nanoparticles, enabling RNA protection and delivery across the cell membrane. (C) in vitro screening of amino acid-derived CARTs and charge-altering dependent mRNA release mechanism. (D) *In vivo* targeted mRNA delivery to lung vasculatures. Figure created with BioRender.com.

## Results and Discussion

### Positioning of the pendent amine group influences *in vitro* and *in vivo* mRNA delivery

To test whether the number of methylenes separating the pendent amino group from the α-carbon influences mRNA delivery efficacy, we initially synthesized ornithine morpholinone monomer (M_orn_) which features a spacer bearing one less methylene than lysine (Figure 1A). Using an optimized organocatalytic ring-opening polymerization (OROP), we prepared the ornithine-derived copolymers (BnO-D_m_-*b*-Orn_n_-OAc, m = 15 or 16, n = 5) by sequential 1,5,7-triazabicyclo[4.4.0]dec-5-ene (TBD)-catalyzed ROP of a dodecyl-functionalized carbonate monomer (D = dodecyl MTC, polymerized at room temperature) to form the lipid block^48^ and M_Orn_ (polymerized at –78 °C) to form the pro-cationic block, followed by end-capping with acetic anhydride. The resulting acetyl-terminated D-Orn-CART_Boc_ copolymer was purified by dialysis and deprotected to yield D-Orn-CART (Figure 1A).

CART/RNA NPs were prepared by pipette mixing of firefly luciferase-encoding (fLuc) mRNA with a DMSO solution of D-Orn-CART at charge ratio of 10 (+/− ratio, the ratio of nitrogen in CART to phosphate in mRNA) in PBS at pH 5.5 (Figure 1B). Physicochemical characterization of the resulting D-Orn-CART/mRNA formulations analyzed by DLS revealed nanoparticles with a Z-average diameter of ∼160-190 nm (Table S1). mRNA encapsulation efficiency of D-Orn-CART/mRNA particles formulated at 10:1, (+/−) charge ratio was measured using a RiboGreen assay and was found to be 96% (Table S1).

We compared the cellular transfection efficiency of the D-Orn-CART to the previously reported D-Gly– and D-Lys-CARTs (Figure 2A) in A549 cells.^48,57^ To investigate both cell entry and functional mRNA delivery, cultured A549 cells were treated with CARTs formulated with Cy5-labeled, eGFP-coding mRNA where GFP signal can be used as a proxy for functional mRNA delivery and Cy5 signal as an indication of cell entry. D-Orn-CART/Cy5-eGFP mRNA NPs formulated at a 10:1 (+/−) charge ratio exhibited comparable levels of protein expression to those treated with N-hydroxyethyllysine-derived CART (Lys-CART, BnO-D_14_-*b*-Lys_8_) as measured by flow cytometry (Figure 2B). On the other hand, EGFP mRNA delivery with N-hydroxylglycine-derived CART (D-Gly-CART) mRNA NPs results in high overall protein expression in A549 cells, showing a greater than fivefold increase in expression relative to D-Orn-CART/mRNA and D-Lys-CART/mRNA (Figure 2B). The higher level of protein expression observed with D-Gly-CART/mRNA NPs in A549 cells relative to D-Orn-CART and D-Lys-CART/mRNA NPs is not attributable to improved cellular uptake. Cells treated with D-Lys-CART show greater Cy5 fluorescence compared to D-Gly CART and D-Orn-CART, indicating that D-Lys-CART/mRNA NPs are internalized more readily by A549 cells (Figure 2C). These data indicate that the variation in performance CART polymers is likely attributable to processes downstream of nanoparticle internalization (for example, endosomal escape, mRNA release or intracellular trafficking).^57^ As previous studies had suggested that the rate of degradation of the cationic component of the CART amphiphiles may be correlated to cellular transfection efficiency,^20^ we investigated the rate at which poly(hydroxyethyl ornithine) (p(HE-Orn)2^+^), the homopolymer of the cationic block of the D-Orn-CART copolymer, rearranges and the products of degradation in buffered water in the absence of an mRNA cargo (Figure S1) and compared that to the degradation of the related glycine and lysine-derived homopolymers.^48,57,65^ When suspended in phosphate-buffered water at pH 6.5, the deprotected homopolymer p(HE-Orn)2^+^ degrades with a t_1/2_ of 16 min to selectively form 3S-(2-hydroxylethylamino)2-piperidinone (O-lactam, Figure S5A, S5B). After 49 minutes, 90% of p(HE-Orn)2^+^ **20** was converted to the lactam; the hydrolysis product N-hydroxyethyl ornithine (HE-Orn)2+ was the only other product identified (<5%, Figures S5A, S5B). In contrast, polycations derived from N-hydroxyethylglycine (Gly-CARTs) degrade by sequential 1,5– and 1,6-O→N acyl shifts to form neutral diketopiperazines (t_1/2_ = 3 min, pH 6.5),^59,66^ polycations derived from N-hydroxyethyllysine degrade more by hydrolysis at a rate (t_1/2_ = 12 min, pH 6.5) comparable to that of the ornithine-derived polycation (Table S2). The high selectivity observed is attributed to the positioning of the pendant side chain amine, which facilitates intramolecular cyclization by a 1,6-O→N acyl shift (Figure S5B). Collectively, these data suggest that fine tuning the balance between complexation/decomplexation (i.e. packaging vs. release) is correlated to the efficiency of functional mRNA delivery in cell culture. While *in vitro* mRNA delivery is a poor predictor of the in vivo efficiency and tropism, these initial results motivated further *in vivo* investigations.

**Figure 2.**
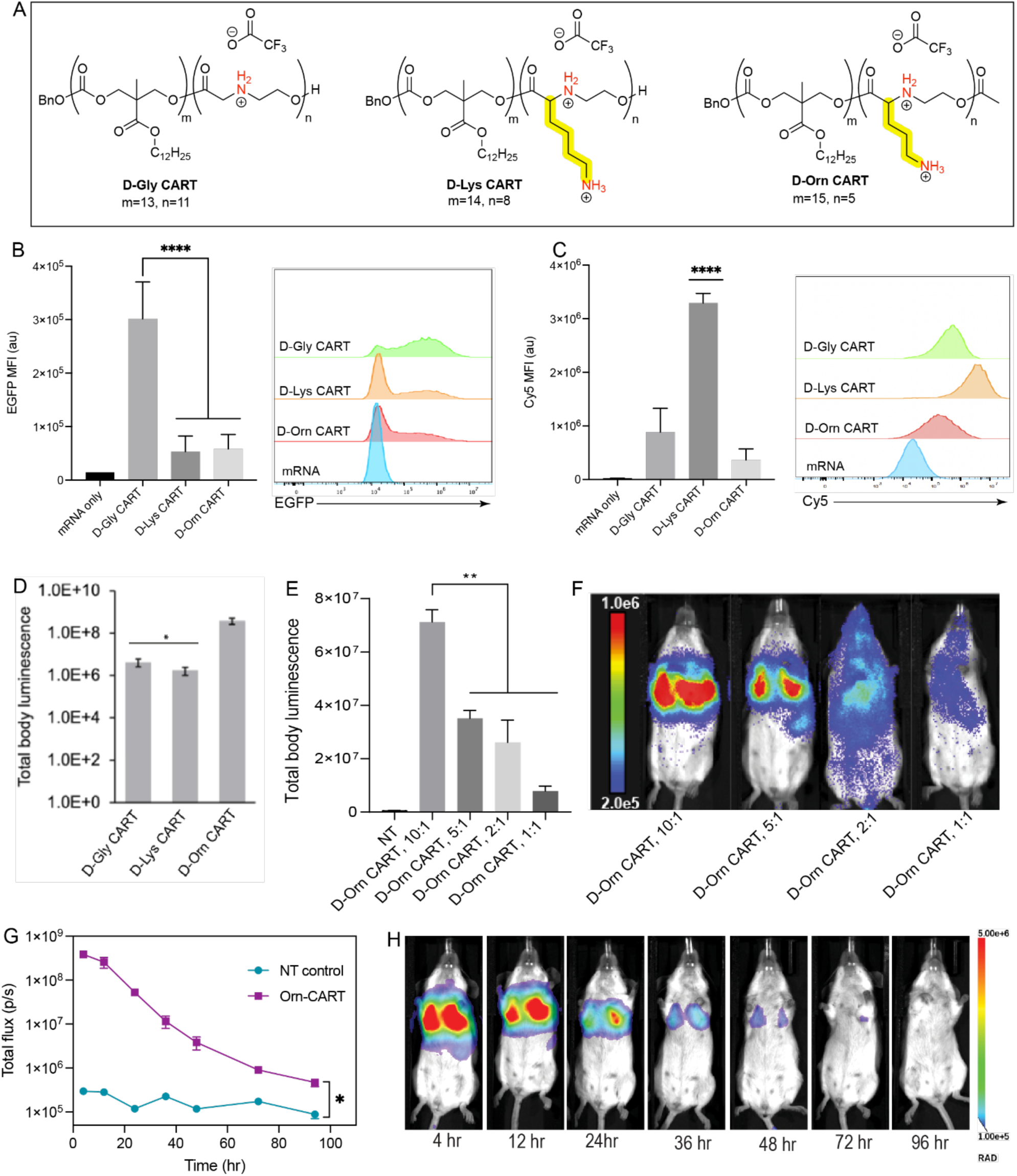
Number and spatial array of cationic charges within the CART polymer dictate both *in vitro* and *in vivo* mRNA delivery efficacy. (A) Chemical structures of D-glycine-, D-lysine-, and D-ornithine-derived CARTs. (B, C) Flow cytometry analysis of A549 cells treated with Cy5-labeled eGFP-encoding mRNA formulated with D-Gly-CART, D-Lys-CART and D-Orn-CART at a 10:1 (+/−) charge ratio (**** *P* < 0.0001. (G) Total body luminescence quantitation of mice injected intravenously with fLuc mRNA formulated at a 10:1 (+/−) charge ratio with D-Gly-CART, D-Lys-CART, or D-Orn-CART, *n* = 3, \**P <* 0.05. (E, F) *In vivo* charge ratio screen of D-Orn-CART and representative bioluminescence images. (G, H) Time course of protein expression in lungs of mice after treatment with D-Orn-CART/fLuc mRNA nanoparticles and representative bioluminescence images of mRNA expression kinetics over 96 hours for mice treated intravenously with 5 μg fLuc mRNA formulated with D-Orn-CART at 10:1 (+/−), n = 3, \**P <* 0.05.

When administered intravenously (tail vein) into mice, D-Orn-CART formulated with firefly luciferase-encoding (fLuc) mRNA at a 10:1 (+/−) charge ratio led to high protein expression localized selectively to the lungs, resulting in total body luminescence more than one order of magnitude greater than that observed with either D-Gly-CART or D-Lys-CART (Figure 2D). As we had previously noted that charge ratio impacts transfection *in vitro*, and because it has been shown that NP charge can impact tissue tropism,^20^ we explored the impact of varying the ratio of D-Orn-CART to mRNA on *in vivo* protein expression. We found that when fLuc mRNA is formulated with D-Orn-CART at a 5:1 (+/−) charge ratio, protein expression shifts; the majority of protein expression is still observed in the lung, but protein expression is now also observed in the spleen. At lower charge ratios (2:1 and 1:1 (+/−)), the distribution profile changes drastically, showing systemic protein expression throughout the mouse (Figure 2E, 2F). To quantify the tissue selectivity of protein expression, D-Orn-CART was formulated with fLuc mRNA at a 10:1 or 5:1 (+/−) charge ratio and administered intravenously to mice. Subsequent isolation and bioluminescence imaging of lungs, spleen, liver, and kidneys revealed that greater than 99% of protein expression was localized to the lungs in mice treated with D-Orn-CART at a 10:1 (+/−) charge ratio (Figure S2A, S2B). Consistent with the results shown in Figure 2E, D-Orn-CART/mRNA particles formulated at a 5:1 (+/−) charge ratio showed protein expression split between the lungs (73%) and the spleen (27%) (Figure S2). This tunable tissue tropism^61^ demonstrates the potential of using a single delivery vector in different formulations to address disease treatments or diagnostics in different organs in an organ-specific way.^61^ Importantly, DLS and CryoEM characterization of D-Orn-CART NP formulated at 10:1 and 5:1 (+/−) charge ratios revealed similar characteristics, suggested that changes in tissue tropism was not driven by physicochemical traits (Figure S3). Similar mRNA delivery efficacy and lung tropism were observed for the Oleyl-Orn-CART compared to the D-Orn-CART suggesting that the nature of the cationic repeat unit of the CART amphiphiles has a more significant influence on tissue tropism (Figure S4) than the nature of the lipid.

Cationic polymers and cationic liposomes are known to target lungs, but have in some cases have been shown to cause inflammation or toxicity.^67–71^ To evaluate the tolerability of mRNA delivery with D-Orn-CART/RNA NPs, markers of liver and kidney toxicity were measured in blood drawn 24 hours after intravenous administration of D-Orn-CART complexed with 5 μg fLuc mRNA. Alanine transferase (AST), aspartate transferase (ALT), and blood urea nitrogen (BUN) levels were not significantly elevated compared to untreated mice suggesting a favorable biosafety profile which could be attributed to the excellent biodegradability of CARTs (Figure S6). To evaluate the chronic toxicity, mice were injected with D-Orn-CART NPs encapsulating fLuc mRNA one dose per week for four weeks. Markers of liver and kidney toxicity were measured in blood drawn 24 hours after the last dose. Results indicated no signs of toxicity (AST, ALT and BUN levels) after repeated dosing (Table S3). Long-lived protein expression was observed for many days after administration of D-Orn-CART which could allow for less frequent dosing in a therapeutic context. Remarkably, bioluminescence signal localized in the lungs remains high (total flux >10^6^ p/s) up to 48 h after treatment, and the signal was still detectable in lungs removed from treated mice 94 h after treatment (Figure 2G, 2H, Figure S7). This expression profile is distinct from our first generation Gly-CARTs expression profile which significantly declined after 24 hours; this finding could be attributed to improved delivery efficacy of D-Orn-CART although the exact molecular mechanism is unclear at this stage.^50^ Collectively, these data suggested that control of spatial array of cationic charges within CART polymers can dictate mRNA delivery efficiency *in vitro* and control the *in vivo* tropism along with the durability of protein expression.

### Positioning of the pendent amine group substantially boosts mRNA delivery efficiency and dictates tissue tropism

Motivated by the impressive behavior of Orn-CART *in vitro* and *in vivo*, we sought to systematically evaluate the impact of varying the carbon chain length of the pendent amino group on the side chain. We further expanded the design space of dicationic monomers by designing a diaminobutyric acid derived monomer (Daba) with two carbons in the pendent amino group and diaminopropionic acid derived monomer (Dapa) with one carbon in the pendent amino group (Figure 3A). D-Daba-CART (BnO-D_18_-*b*-Daba_5_-Ac) and D-Dapa-CART (BnO-D_15_-*b*-Dapa_5_-Ac) were prepared following a similar ROP synthetic procedure to that used to make D-Orn-CART. With a series of CARTs varying in the number of carbons in the pendent chain, we then carried out bioluminescence quantification of mice treated intravenously with 5 μg fLuc mRNA formulated individually with D-Lys-CART, D-Orn-CART, D-Daba-CART, or D-Dapa-CART at a 10:1 (+/−) charge ratio (Figure 3B). We also benchmarked lung-targeting CARTs against a commercially available vehicle for lung transfection, by treating mice with 5 μg fLuc mRNA formulated with in vivo-jet PEI®. D-Dapa-CART showed the highest levels of bioluminescent expression followed closely by D-Daba-CART. These CARTs exhibit a total flux around 1×10^9^ (p/s), half an order of magnitude better than D-Orn-CART (5×10^8^ (p/s)), which performed an order of magnitude better than Lys-CART. In vivo-JetPEI performed the worst. Notably, the D-Daba-CART, similar to the D-Lys and D-Orn CARTs, demonstrated highly selective protein expression in the lung whereas the D-Dapa-CART delivered fLuc mRNA indiscriminately to the lungs, liver, and spleen (Figure 3C).

**Figure 3.**
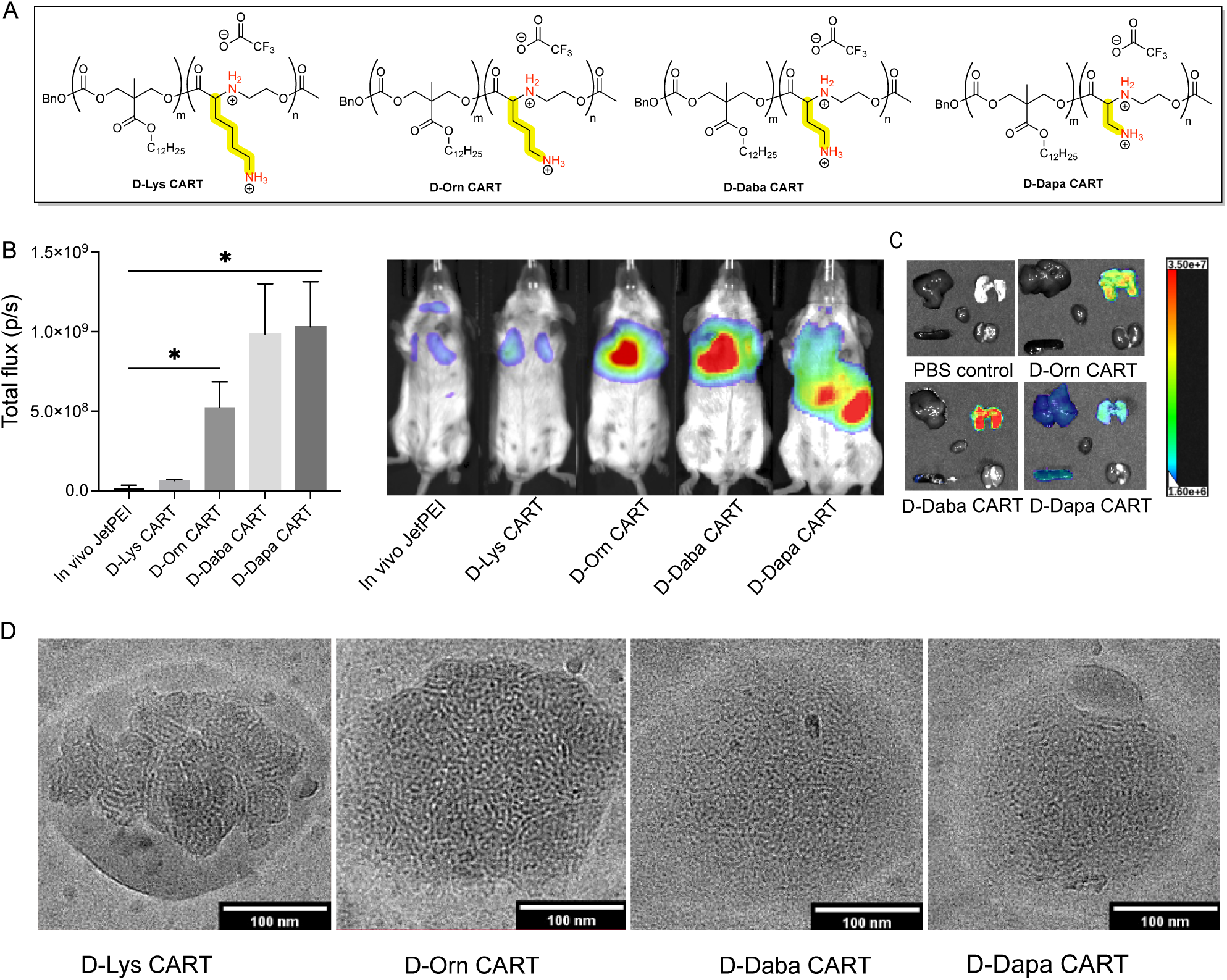
Systematic engineering of the sidechain chemistry of CARTs for enhanced lung delivery and controlled *in vivo* tropism. (A) Design and chemical structures of morpholinone monomers with different spacing between amino groups. (B) Full body bioluminescence quantitation of mice treated intravenously with 5 μg fLuc mRNA formulated with *in vivo*Jet PEI, D-Lys-CART, D-Orn-CART, D-Daba-CART, D-Dapa-CART and representative bioluminescence images (n = 3, *\*P <* 0.05). (C) Representative bioluminescence images of excised organs of mice treated intravenously with 5 μg fLuc mRNA formulated with different dicationic CARTs. (D) CryoEM images of dication CARTs formulated with fLuc mRNA at a 10:1 (+/−) charge ratio.

The morphological and physicochemical characteristics of the dicationic CARTs series were investigated by several techniques. By DLS, all CART/mRNA complexes were similar in size (∼180 nm, Table S1). CryoEM showed the presence of roughly spherical nanoparticles with an internal disordered bicontinuous morphology with a periodicity of about 6 nm.^72^ D-Dapa-CART had the highest zeta potential (+82 mV) followed by D-Daba and D-Orn-CART (+73 mV), while D-Lys-CART had the lowest zeta potential (+52 mV). Small-angle X-ray scattering (SAXS) further confirmed this feature with a broad scattering peak centered q ≈ 0.1 Å^-1^ corresponding to spacings of the bicontinuous domains averaging 6.2 nm (Figure S8).^73–74^ The D-Dapa CART/RNA NP, the only non-lung selective CART in this series, exhibited blebs in the CryoEM images (Figure 3D) and a SAXS scattering feature slightly shifted to higher q (q ≈ 0.11 Å^-1^, corresponding to a d-spacing of approx. 5.8 nm).^75^ Further studies are underway to assess if these morphological differences are correlated with transfection efficiency and organ selectivity.

### Lung-selective CARTs target pulmonary endothelium with high specificity

An understanding of functional RNA delivery at a cellular level can be enabling for development of targeted therapies for lung disease. Toward this end, we further quantitatively analyzed functional mRNA delivery on a single cell level by determining the specific cell populations within the lung that express protein upon treatment with D-Orn-CART. D-Orn-CART was formulated with 10 ug NanoLuc-encoding mRNA at a 10:1 (+/−) charge ratio and administered intravenously. Lung single cell suspensions from treated mice were sorted by FACS into epithelial, endothelial, and immune cell populations (10,000 cells per population). Luminescence in each cellular subpopulation was measured after addition of Nano-Glo substrate to quantify protein expression. We found that treatment with D-Orn-CART/mRNA NPs at a 10:1 (+/−) charge ratio results in protein expression primarily in endothelial cells in the lung (Figures S9 and S10). Endothelial cells are highly affected in a variety of respiratory diseases and are therefore a promising target for therapeutic gene delivery.^76^ However, the vascular bed of the lung is complex and composed of at least six major endothelial cell types (pulmonary artery, pulmonary vein, gCap and aerocyte capillaries, bronchial vessels, and lymphatics) with distinct functions and precise spatial positioning.^77–78^ ^78–79^ To determine the identity of endothelial subtypes within the lung that were targeted by D-Orn-CART and D-Daba-CART, and their location and uptake efficiency, we turned to quantitative histological analysis. To indelibly mark any cells in which CART mRNA contents had been delivered and translated into functional protein, we packaged Cre recombinase mRNA using either D-Orn-CART, D-Daba-CART or D-Lys-CART and injected them individually into Ai14 Cre reporter mice in which cells are marked by TdTomato fluorescent protein expression following Cre-mediated recombination (Figure 4A).^80^ Kidney, heart, spleen, liver, and lungs were collected and examined for TdTomato expression. While scattered tdTomato-marked cells were present in kidney, heart, spleen, and liver (Figure S11) the overwhelming majority of TdTomato fluorescence was found in the lung as expected from our prior studies (Fig 4B-D). Detailed analysis of lung targeting was performed on D-Orn-CART and D-Daba-CART animals, as D-Lys-CART demonstrated significantly lower recombination in the pulmonary vasculature (Figure S12). In the lungs of both D-Orn-CART and D-Daba-CART animals abundant TdTomato expression was apparent in the lining of pulmonary arteries and veins and within the alveoli, but no TdTomato was visible either in the conducting airway epithelium or in smooth muscle cells surrounding either airways or vessels (Figure 4E&F). High resolution confocal imaging of the pulmonary circulation revealed that in both pulmonary arteries and veins TdTomato labeling was confined to the endothelium (217 of 217 and 862 of 862 TdTomato+ cells express the endothelial marker CD31 in D-Orn-CART and D-Daba-CART; respectively (Figure 5). The complex cellular architecture, extremely flattened morphology and close apposition of epithelium and endothelium within alveoli makes capillary cell type identification using cell surface proteins such as CD31 unreliable. Therefore, an antibody to the endothelial-specific ETS-related gene (ERG) was used to unambiguously identify alveolar capillary nuclei in conjunction with tdTomato signal. Using this approach, we found that the majority of TdTomato-marked cells were ERG+ in both D-Orn-CART and D-Daba-CART (417 of 438 and 525 of 581 TdTomato+ cells express ERG in D-Orn-CART and D-Daba-CART, respectively), indicating that non-capillary alveolar cells are minimally labeled by this method. Collectively, these findings indicate that D-Orn-CART and D-Daba-CART target the lung and lung endothelium with a high degree of selectivity.

**Figure 4:**
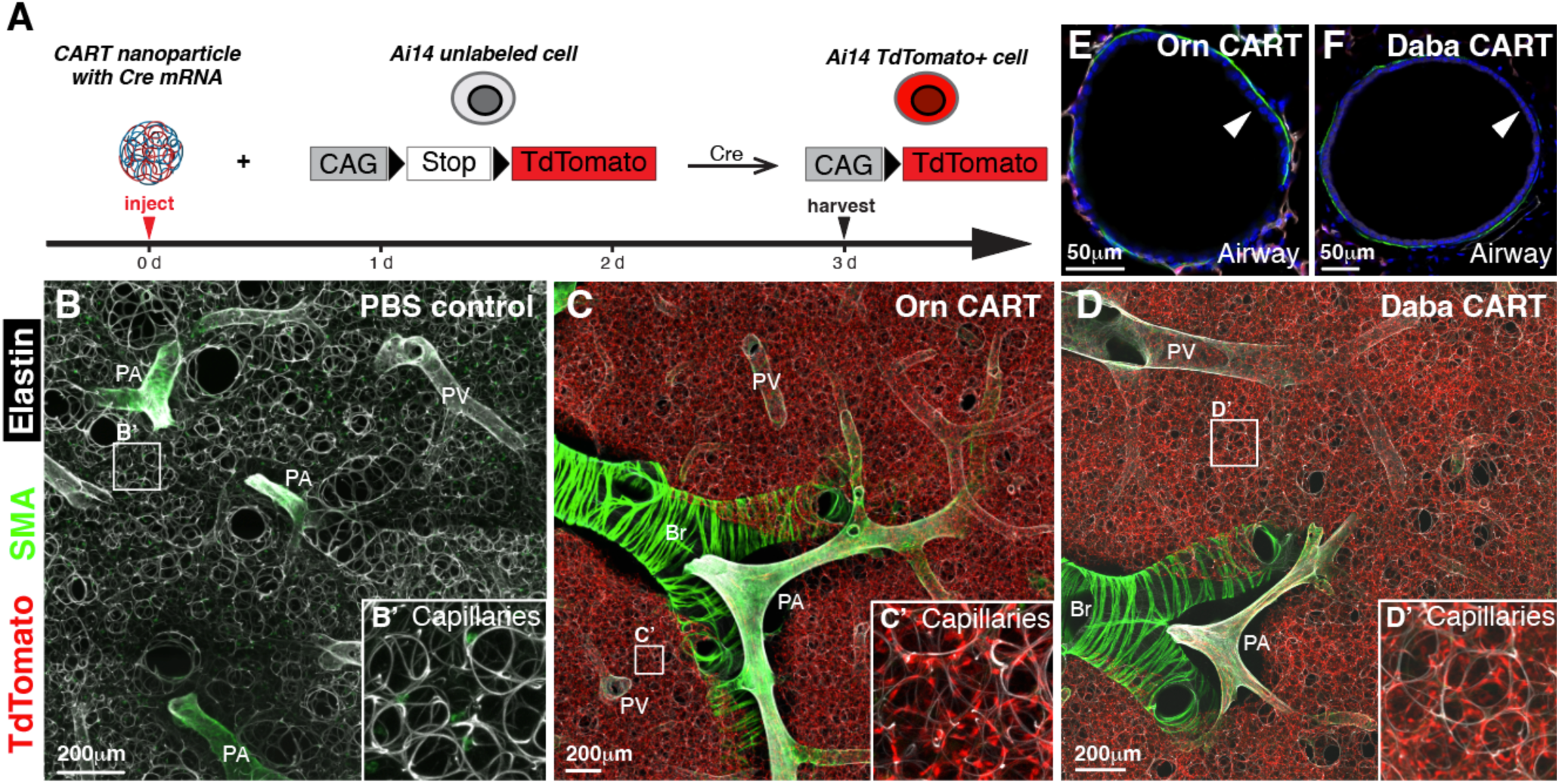
Cre mRNA delivery using D-Orn– and D-Daba-CARTs induces robust Cre reporter recombination in the pulmonary endothelium. (A) Schematic of CART, Ai14 Cre reporter mice and experimental strategy used to test RNA delivery in vivo. (B) PBS-injected Ai14 Cre reporter mouse. (C) D-Orn-CART delivery of Cre mRNA into Ai14 Cre reporter mouse. (D) D-Daba-CART delivery of Cre mRNA into Ai14 Cre reporter mouse. Airway epithelium is unlabeled in both D-Orn-CART (E) and D-Daba-CART (F). (B-D) Maximum intensity projections of tiled confocal z stacks of 300mm vibratome sections stained to highlight smooth muscle a-actin (SMA, green) and elastin (white); TdTomato direct fluorescence from recombined Ai14 Cre reporter (red). No targeting of smooth muscle cells of conducting airways is observed. Insets B’-D’, show alveolar capillary detail. (E, F), Cryosections stained to highlight smooth muscle (green), TdTomato from recombined Ai14 Cre reporter mouse (red), CD31 (white), and DAPI nuclei (blue). PA, pulmonary artery; PV, pulmonary vein; Br, airway bronchus.

**Figure 5:**
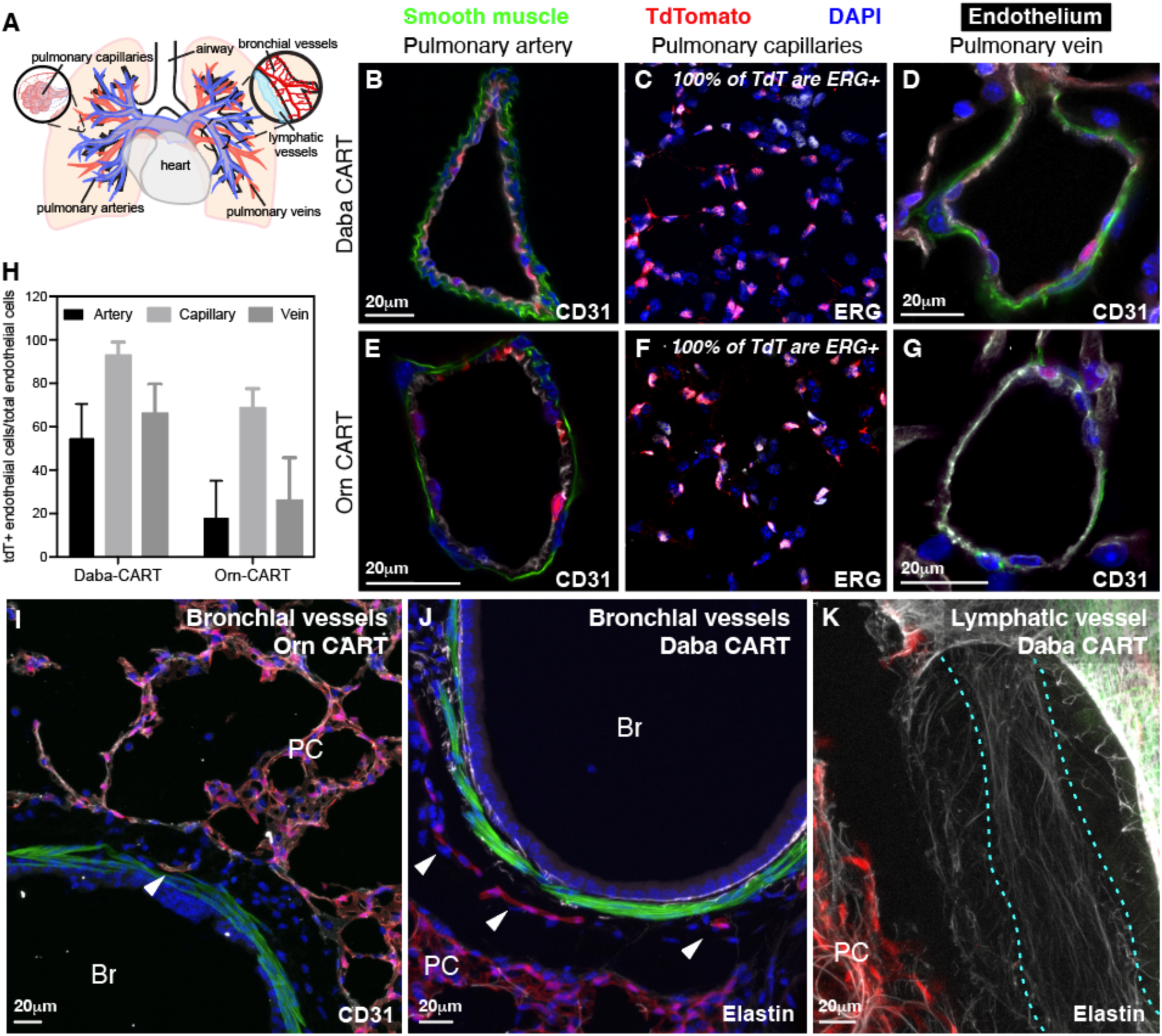
D-Orn-CART and D-Daba-CART exclusively target endothelial cells in the pulmonary circulation. (A) Schematic of diversity and distribution of lung endothelia. (B-D) Endothelium of pulmonary artery (B), capillaries (C), and veins (D) and not smooth muscle cells are TdTomato+ following Cre mRNA delivery by D-Daba-CART into Ai14 Cre reporter mouse. (E-G) Endothelium of pulmonary artery (E), capillaries (F), and veins (G) and not smooth muscle cells are TdTomato+ following Cre mRNA delivery by D-Orn-CART into Ai14 Cre reporter mouse. (H) Percent of endothelial cells marked by TdTomato fluorescence in each compartment of the pulmonary circulation. Scattered TdTomato labeling among bronchial vessels (arrowheads) in D-Orn-CART (I) and D-Daba-CART (J, K), in all cases lymphatic endothelium (lymphatic vessel outlined with cyan dotted line) was unlabeled, D-Daba-CART shown. (B-G) & (I-K) Cryosections stained to highlight smooth muscle a-actin (SMA, green), TdTomato direct fluorescence from recombined Ai14 Cre reporter (red), and DAPI nuclei (blue). White, endothelium shown in B-I (B, D, E, G & I, CD31; C&F, ERG), elastin shown in J & K. (B-G) single confocal optical sections. (I-K) maximum intensity projections of 20μm confocal z stacks. Br, airway bronchus; PC, pulmonary capillaries. *n* = 3 biologically independent animals.

### The top lung-selective CARTs target pulmonary arteries, capillaries and veins with high efficiency

The artery endothelium was efficiently labeled by D-Daba-CART (53% artery endothelia TdTomato+; 956 endothelial cells from 61 arteries counted; Fig 5B and 5H) while the D-Orn-CART showed lower efficiency (26% artery endothelial cells are tdTomato+; 465 endothelial cells from 55 arteries counted; Figure 5E and 5H). For the D-Daba-CART, veins demonstrated significantly higher recombination rates than arteries (65% vein endothelial cells are tdTomato+ in D-Daba-CART, 542 endothelial cells from 66 veins counted, p<0.01) (Figure 5D and 5H), but not in D-Orn-CART (31% vein endothelial cells are tdTomato+ in D-Orn-CART, 311 endothelial cells from 48 veins counted, p=0.16; Figure 5G and 5H). D-Daba-CART exhibited exceptionally efficient targeting of the pulmonary capillaries, with 93% of capillary cells expressing tdTomato (525 tdTomato+ of 564 ERG+ cells counted, Figure 5C and 5H), indicating targeting of both aerocytes and gCaps, the two capillary cell types. Efficiency was lower for the D-Orn-CART with 66% of capillary cells expressing tdTomato (417 tdTomato+ of 632 ERG+ cells counted; Figure 5F and 5H). We also observed some variability in cell targeting efficiency between individual animals, both within the D-Orn-CART and D-Daba-CART treatment groups (Figure S13). Capillary targeting was significantly higher than that in pulmonary artery and pulmonary vein endothelium in both D-Daba– and D-Orn-CART (p<0.01 in all cases). Neither D-Daba-CART nor D-Orn-CART targeted lymphatic endothelium and both demonstrated low efficiency in targeting bronchial arteries (Figure 5I-K).

## Conclusion

Here we show that a series of dicationic amino acid derived CARTs is highly effective for delivery of mRNA to the lung vasculature after systemic administration in mice. Importantly, we show that systematic variation of the side chain of the CART cationic block has a significant influence on not only mRNA delivery efficiency but also on the locus of protein expression upon systemic administration of CART/RNA nanoparticles (NPs) in mice. The remarkable sensitivity of mRNA delivery efficacy to subtle perturbations in transporter structure motivates further investigation into the features of polymers that enable effective and tissue selective RNA delivery.^81^ Furthermore, the ability to tune the locus of gene expression in vivo by modifying CART polymer structure lends support to the hypothesis that tissue tropism can be controlled by intrinsic nanoparticle properties without the need for targeting ligands, presenting opportunities for development of simple and efficient gene therapies. The efficient synthesis of this new family of CART amphiphiles, coupled with their selective and tunable lung endothelium targeting without molecular ligands, represents an enabling platform for research and clinical applications. Overall, the lung selective CARTs developed in this study are anticipated to accelerate the clinical development of mRNA therapeutic approaches to address unmet medical needs in pulmonary vascular diseases by providing delivery tools and fundamental knowledge to advance lung targeted gene therapies. Continued development of effective CARTs to deliver nucleic acids to lung-relevant cell types *in vivo* may eventually enable a new therapy for vascular endothelial lung diseases.

## Author contribution

M.M.A, S.R.B., J.N., R.L.M., R.L., P.A.W., M.E.K. and R.M.W designed research; M.M.A., J.N, S.R.M, M.E.K., S.R.B., R.L.M., P.J.H., S.R.K, T.R.B., Y.J., A.S., O.A.W.H., I.S., D.K.C. performed experiments; M.M.A, J.N., S.R.B., R.L.M., P.J.H., and R.M.W analyzed data; M.M.A, J.N., S.R.B., R.L.M., P.J.H., M.E.K., P.A.W. and R.M.W wrote the paper.

## Acknowledgements

This work was supported by grants 5R01CA245533-03 (R.M.W., P.A.W., R.L.), R01 CA031845 and R01 AI161803 (P.A.W.) and R01HL163013 (M.E.K.) from the National Institutes of Health, and grants CHE-2002933 (R.M.W.) from the National Science Foundation, the Child Health Research Institute at Stanford University, and the SPARK Translational Research Program in the Stanford University School of Medicine (R.M.W., P.A.W., R.L.). Support through the Stanford Cancer Translational Nanotechnology Training T32 Training Grant T32 CA196585 funded by the National Cancer Institute (T.R.B), the Stanford Training Program in Lung Biology 5T32HL129970-08 funded by the National Heart, Lung and Blood Institute (J.N), through the Stanford Maternal & Child Health Research Institute (J.N), through the Center for Molecular Analysis and Design at Stanford University (R.L.M), and through the National Science Foundation Graduate Research Fellowships Program (DGE-1656518, S.R.B.) is also acknowledged. Some of the cell sorting/flow cytometry analysis for this project was done on instruments in the Stanford Shared FACS Facility. The authors thank VLP Therapeutics for providing NanoLuc-encoding mRNA. This research used the resources of the Stanford cryoEM center, notably the Glacios cryoEM instrument. We would like to thank staff scientists Dr. Bharti Signal for training and assistance. We would like to thank staff scientist Dr. Greg Hura for the collection of the SAXS data acquired at the Advanced Light Source (ALS), a national user facility operated by Lawrence Berkeley National Laboratory on behalf of the Department of Energy, Office of Basic Energy Sciences, through the Integrated Diffraction Analysis Technologies (IDAT) program, supported by DOE Office of Biological and Environmental Research. Additional support comes from the National Institute of Health project ALS-ENABLE (P30 GM124169) and a High-End Instrumentation Grant S10OD018483.

## References

1. Parhiz, H.; Atochina-Vasserman, E. N.; Weissman, D., mRNA-based therapeutics: looking beyond COVID-19 vaccines. Lancet 2024, 403 (10432), 1192–1204.

2. Berger, S.; Lachelt, U.; Wagner, E., Dynamic carriers for therapeutic RNA delivery. Proc Natl Acad Sci U S A 2024, 121 (11), e2307799120.

3. Cullis, P. R.; Felgner, P. L., The 60-year evolution of lipid nanoparticles for nucleic acid delivery. Nature Reviews Drug Discovery 2024, 23 (9), 709–722.

4. Verbeke, R.; Lentacker, I.; De Smedt, S. C.; Dewitte, H., The dawn of mRNA vaccines: The COVID-19 case. Journal of Controlled Release 2021, 333, 511–520.

5. Kumar, R.; Santa Chalarca, C. F.; Bockman, M. R.; Bruggen, C. V.; Grimme, C. J.; Dalal, R. J.; Hanson, M. G.; Hexum, J. K.; Reineke, T. M., Polymeric Delivery of Therapeutic Nucleic Acids. Chem. Rev. 2021, 121 (18), 11527–11652.

6. Reichmuth, A. M.; Oberli, M. A.; Jaklenec, A.; Langer, R.; Blankschtein, D., mRNA vaccine delivery using lipid nanoparticles. Ther Deliv 2016, 7 (5), 319–334.

7. Dilliard, S. A.; Siegwart, D. J., Passive, active and endogenous organ-targeted lipid and polymer nanoparticles for delivery of genetic drugs. Nature Reviews Materials 2023, 8 (4), 282–300.

8. Kim, J.; Eygeris, Y.; Ryals, R. C.; Jozić, A.; Sahay, G., Strategies for non-viral vectors targeting organs beyond the liver. Nature Nanotechnology 2024, 19 (4), 428–447.

9. Jain, M.; Yu, X.; Schneck, J. P.; Green, J. J., Nanoparticle Targeting Strategies for Lipid and Polymer-Based Gene Delivery to Immune Cells In Vivo. Small Sci 2024, 4 (9), 2400248.

10. Piotrowski-Daspit, A. S.; Bracaglia, L. G.; Eaton, D. A.; Richfield, O.; Binns, T. C.; Albert, C.; Gould, J.; Mortlock, R. D.; Egan, M. E.; Pober, J. S.; Saltzman, W. M., Enhancing in vivo cell and tissue targeting by modulation of polymer nanoparticles and macrophage decoys. Nat Commun 2024, 15 (1), 4247.

11. Litzinger, D. C.; Buiting, A. M. J.; van Rooijen, N.; Huang, L., Effect of liposome size on the circulation time and intraorgan distribution of amphipathic poly(ethylene glycol)-containing liposomes. Biochim. Biophys. Acta 1994, 1190 (1), 99–107.

12. Caster, J. M.; Yu, S. K.; Patel, A. N.; Newman, N. J.; Lee, Z. J.; Warner, S. B.; Wagner, K. T.; Roche, K. C.; Tian, X.; Min, Y.; Wang, A. Z., Effect of particle size on the biodistribution, toxicity, and efficacy of drug-loaded polymeric nanoparticles in chemoradiotherapy. Nanomedicine 2017, 13 (5), 1673–1683.

13. Kaga, S.; Truong, N. P.; Esser, L.; Senyschyn, D.; Sanyal, A.; Sanyal, R.; Quinn, J. F.; Davis, T. P.; Kaminskas, L. M.; Whittaker, M. R., Influence of Size and Shape on the Biodistribution of Nanoparticles Prepared by Polymerization-Induced Self-Assembly. Biomacromolecules 2017, 18 (12), 3963–3970.

14. He, C.; Hu, Y.; Yin, L.; Tang, C.; Yin, C., Effects of particle size and surface charge on cellular uptake and biodistribution of polymeric nanoparticles. Biomaterials 2010, 31 (13), 3657–3666.

15. Elci, S. G.; Jiang, Y.; Yan, B.; Kim, S. T.; Saha, K.; Moyano, D. F.; Yesilbag Tonga, G.; Jackson, L. C.; Rotello, V. M.; Vachet, R. W., Surface Charge Controls the Suborgan Biodistributions of Gold Nanoparticles. ACS Nano 2016, 10 (5), 5536–5542.

16. Ling, D.; Hackett, M. J.; Hyeon, T., Surface ligands in synthesis, modification, assembly and biomedical applications of nanoparticles. Nano Today 2014, 9 (4), 457–477.

17. Brown, S. B.; Wang, L.; Jungels, R. R.; Sharma, B., Effects of cartilage-targeting moieties on nanoparticle biodistribution in healthy and osteoarthritic joints. Acta Biomaterialia 2020, 101, 469–483.

18. Monopoli, M. P.; Åberg, C.; Salvati, A.; Dawson, K. A., Biomolecular coronas provide the biological identity of nanosized materials. Nature Nanotechnology 2012, 7 (12), 779–786.

19. Venkataraman, S.; Hedrick, J. L.; Ong, Z. Y.; Yang, C.; Ee, P. L. R.; Hammond, P. T.; Yang, Y. Y., The effects of polymeric nanostructure shape on drug delivery. Advanced Drug Delivery Reviews 2011, 63 (14), 1228–1246.

20. Kranz, L. M.; Diken, M.; Haas, H.; Kreiter, S.; Loquai, C.; Reuter, K. C.; Meng, M.; Fritz, D.; Vascotto, F.; Hefesha, H.; Grunwitz, C.; Vormehr, M.; Hüsemann, Y.; Selmi, A.; Kuhn, A. N.; Buck, J.; Derhovanessian, E.; Rae, R.; Attig, S.; Diekmann, J.; Jabulowsky, R. A.; Heesch, S.; Hassel, J.; Langguth, P.; Grabbe, S.; Huber, C.; Türeci, Ö.; Sahin, U., Systemic RNA delivery to dendritic cells exploits antiviral defence for cancer immunotherapy. Nature 2016, 534 (7607), 396–401.

21. Fenton, O. S.; Kauffman, K. J.; Kaczmarek, J. C.; McClellan, R. L.; Jhunjhunwala, S.; Tibbitt, M. W.; Zeng, M. D.; Appel, E. A.; Dorkin, J. R.; Mir, F. F.; Yang, J. H.; Oberli, M. A.; Heartlein, M. W.; DeRosa, F.; Langer, R.; Anderson, D. G., Synthesis and Biological Evaluation of Ionizable Lipid Materials for the In Vivo Delivery of Messenger RNA to B Lymphocytes. Adv. Mater. 2017, 29 (33), 1606944.

22. Meyer, R. A.; Neshat, S. Y.; Green, J. J.; Santos, J. L.; Tuesca, A. D., Targeting strategies for mRNA delivery. Materials Today Advances 2022, 14, 100240.

23. Li, Z.; Amaya, L.; Pi, R.; Wang, S. K.; Ranjan, A.; Waymouth, R. M.; Blish, C. A.; Chang, H. Y.; Wender, P. A., Charge-altering releasable transporters enhance mRNA delivery in vitro and exhibit in vivo tropism. Nat. Commun. 2023, 14 (1).

24. Meshanni, J. A.; Stevenson, E. R.; Zhang, D.; Sun, R.; Ona, N. A.; Reagan, E. K.; Abramova, E.; Guo, C. J.; Wilkinson, M.; Baboo, I.; Yang, Y.; Pan, L.; Maurya, D. S.; Percec, V.; Li, Y.; Gow, A.; Weissman, D.; Atochina-Vasserman, E. N., Targeted delivery of TGF-beta mRNA to murine lung parenchyma using one-component ionizable amphiphilic Janus Dendrimers. Nat Commun 2025, 16 (1), 1806.

25. Yazdi, M.; Pohmerer, J.; Hasanzadeh Kafshgari, M.; Seidl, J.; Grau, M.; Hohn, M.; Vetter, V.; Hoch, C. C.; Wollenberg, B.; Multhoff, G.; Bashiri Dezfouli, A.; Wagner, E., In Vivo Endothelial Cell Gene Silencing by siRNA-LNPs Tuned with Lipoamino Bundle Chemical and Ligand Targeting. Small 2024, 20 (42), e2400643.

26. Qiu, M.; Tang, Y.; Chen, J.; Muriph, R.; Ye, Z.; Huang, C.; Evans, J.; Henske, E. P.; Xu, Q., Lung-selective mRNA delivery of synthetic lipid nanoparticles for the treatment of pulmonary lymphangioleiomyomatosis. Proc Natl Acad Sci USA 2022, 119 (8), e2116271119.

27. Jarzębińska, A.; Pasewald, T.; Lambrecht, J.; Mykhaylyk, O.; Kümmerling, L.; Beck, P.; Hasenpusch, G.; Rudolph, C.; Plank, C.; Dohmen, C., A Single Methylene Group in Oligoalkylamine-Based Cationic Polymers and Lipids Promotes Enhanced mRNA Delivery. Angew. Chem. Int. Ed. 2016, 55, 9591.

28. Kaczmarek, J. C.; Kauffman, K. J.; Fenton, O. S.; Sadtler, K.; Patel, A. K.; Heartlein, M. W.; DeRosa, F.; Anderson, D. G., Optimization of a Degradable Polymer–Lipid Nanoparticle for Potent Systemic Delivery of mRNA to the Lung Endothelium and Immune Cells. Nano Letters 2018, 18 (10), 6449–6454.

29. Kaczmarek, J. C.; Patel, A. K.; Kauffman, K. J.; Fenton, O. S.; Webber, M. J.; Heartlein, M. W.; DeRosa, F.; Anderson, D. G., Polymer–Lipid Nanoparticles for Systemic Delivery of mRNA to the Lungs. Angew. Chem. Int. Ed. 2016, 55 (44), 13808–13812.

30. Schrom, E.; Huber, M.; Aneja, M.; Dohmen, C.; Emrich, D.; Geiger, J.; Hasenpusch, G.; Herrmann-Janson, A.; Kretzschmann, V.; Mykhailyk, O.; Pasewald, T.; Oak, P.; Hilgendorff, A.; Wohlleber, D.; Hoymann, H.-G.; Schaudien, D.; Plank, C.; Rudolph, C.; Kubisch-Dohmen, R., Translation of Angiotensin-Converting Enzyme 2 upon Liver– and Lung-Targeted Delivery of Optimized Chemically Modified mRNA. Molecular Therapy – Nucleic Acids 2017, 7, 350–365.

31. Yan, Y.; Xiong, H.; Zhang, X.; Cheng, Q.; Siegwart, D. J., Systemic mRNA Delivery to the Lungs by Functional Polyester-based Carriers. Biomacromolecules 2017, 18 (12), 4307–4315.

32. Kowalski, P. S.; Capasso Palmiero, U.; Huang, Y.; Rudra, A.; Langer, R.; Anderson, D. G., Ionizable Amino-Polyesters Synthesized via Ring Opening Polymerization of Tertiary Amino-Alcohols for Tissue Selective mRNA Delivery. Adv. Mater. 2018, 30 (34), 1801151.

33. Li, Q.; Chan, C.; Peterson, N.; Hanna, R. N.; Alfaro, A.; Allen, K. L.; Wu, H.; Dall’Acqua, W. F.; Borrok, M. J.; Santos, J. L., Engineering Caveolae-Targeted Lipid Nanoparticles To Deliver mRNA to the Lungs. ACS Chem. Bio. 2020, 15 (4), 830–836.

34. Abd Elwakil, M. M.; Gao, T.; Isono, T.; Sato, Y.; Elewa, Y. H. A.; Satoh, T.; Harashima, H., Engineered ε-decalactone lipomers bypass the liver to selectively in vivo deliver mRNA to the lungs without targeting ligands. Materials Horizons 2021, 8 (8), 2251–2259.

35. Liu, S.; Cheng, Q.; Wei, T.; Yu, X.; Johnson, L. T.; Farbiak, L.; Siegwart, D. J., Membrane-destabilizing ionizable phospholipids for organ-selective mRNA delivery and CRISPR–Cas gene editing. Nat. Mater. 2021, 20, 701–710.

36. Zhang, D.; Atochina-Vasserman, E. N.; Maurya, D. S.; Huang, N.; Xiao, Q.; Ona, N.; Liu, M.; Shahnawaz, H.; Ni, H.; Kim, K.; Billingsley, M. M.; Pochan, D. J.; Mitchell, M. J.; Weissman, D.; Percec, V., One-Component Multifunctional Sequence-Defined Ionizable Amphiphilic Janus Dendrimer Delivery Systems for mRNA. Journal of the American Chemical Society 2021, 143 (31), 12315–12327.

37. Park, Y.; Moses, A. S.; Demessie, A. A.; Singh, P.; Lee, H.; Korzun, T.; Taratula, O. R.; Alani, A. W. G.; Taratula, O., Poly(aspartic acid)-Based Polymeric Nanoparticle for Local and Systemic mRNA Delivery. Molecular Pharmaceutics 2022, 19 (12), 4696–4704.

38. Petersen, D. M. S.; Weiss, R. M.; Hajj, K. A.; Yerneni, S. S.; Chaudhary, N.; Newby, A. N.; Arral, M. L.; Whitehead, K. A., Branched-Tail Lipid Nanoparticles for Intravenous mRNA Delivery to Lung Immune, Endothelial, and Alveolar Cells in Mice. Adv Healthc Mater 2024, 13 (22), e2400225.

39. Zamora, M. E.; Omo-Lamai, S.; Patel, M. N.; Wu, J.; Arguiri, E.; Muzykantov, V. R.; Myerson, J. W.; Marcos-Contreras, O. A.; Brenner, J. S., Combination of Physicochemical Tropism and Affinity Moiety Targeting of Lipid Nanoparticles Enhances Organ Targeting. Nano Letters 2024, 24 (16), 4774–4784.

40. Sun, Y.; Chatterjee, S.; Lian, X.; Traylor, Z.; Sattiraju, S. R.; Xiao, Y.; Dilliard, S. A.; Sung, Y.-C.; Kim, M.; Lee, S. M.; Moore, S.; Wang, X.; Zhang, D.; Wu, S.; Basak, P.; Wang, J.; Liu, J.; Mann, R. J.; LePage, D. F.; Jiang, W.; Abid, S.; Hennig, M.; Martinez, A.; Wustman, B. A.; Lockhart, D. J.; Jain, R.; Conlon, R. A.; Drumm, M. L.; Hodges, C. A.; Siegwart, D. J., In vivo editing of lung stem cells for durable gene correction in mice. Science 2024, 384 (6701), 1196–1202.

41. Liu, B.; Sajiki, Y.; Littlefield, N.; Hu, Y.; Stuart, W. D.; Sridharan, A.; Cui, X.; Siefert, M. E.; Araki, K.; Ziady, A. G.; Shi, D.; Whitsett, J. A.; Maeda, Y., PBAE-PEG-based lipid nanoparticles for lung cell-specific gene delivery. Molecular Therapy 2025, 33 (3), 1154–1165.

42. Bals, R.; Hiemstra, P. S., Innate immunity in the lung: how epithelial cells fight against respiratory pathogens. European Respiratory Journal 2004, 23 (2), 327.

43. Asha, K.; Kumar, P.; Sanicas, M.; Meseko, C. A.; Khanna, M.; Kumar, B., Advancements in Nucleic Acid Based Therapeutics against Respiratory Viral Infections. Journal of Clinical Medicine 2019, 8 (1), 6.

44. Grumelli, S.; Corry, D. B.; Song, L.-Z.; Song, L.; Green, L.; Huh, J.; Hacken, J.; Espada, R.; Bag, R.; Lewis, D. E.; Kheradmand, F., An Immune Basis for Lung Parenchymal Destruction in Chronic Obstructive Pulmonary Disease and Emphysema. PLOS Medicine 2004, 1 (1), e8.

45. Kim, E. Y.; Battaile, J. T.; Patel, A. C.; You, Y.; Agapov, E.; Grayson, M. H.; Benoit, L. A.; Byers, D. E.; Alevy, Y.; Tucker, J.; Swanson, S.; Tidwell, R.; Tyner, J. W.; Morton, J. D.; Castro, M.; Polineni, D.; Patterson, G. A.; Schwendener, R. A.; Allard, J. D.; Peltz, G.; Holtzman, M. J., Persistent activation of an innate immune response translates respiratory viral infection into chronic lung disease. Nature Medicine 2008, 14 (6), 633–640.

46. Griesenbach, U.; Alton, E. W. F. W., Moving forward: cystic fibrosis gene therapy. Human Molecular Genetics 2013, 22 (R1), R52–R58.

47. Keil, T. W. M.; Baldassi, D.; Merkel, O. M., T-cell targeted pulmonary siRNA delivery for the treatment of asthma. WIREs Nanomedicine and Nanobiotechnology 2020, 12 (5), e1634.

48. McKinlay, C. J.; Vargas, J. R.; Blake, T. R.; Hardy, J. W.; Kanada, M.; Contag, C. H.; Wender, P. A.; Waymouth, R. M., Charge-altering releasable transporters (CARTs) for the delivery and release of mRNA in living animals. Proc Natl Acad Sci USA 2017, 114, E448–E456.

49. Benner, N. L.; Near, K. E.; Bachmann, M. H.; Contag, C. H.; Waymouth, R. M.; Wender, P. A., Functional DNA Delivery Enabled by Lipid-Modified Charge-Altering Releasable Transporters (CARTs). Biomacromolecules 2018, 19, 2812–2824.

50. Haabeth, O. A. W.; Blake, T. R.; McKinlay, C. J.; Waymouth, R. M.; Wender, P. A.; Levy, R., mRNA vaccination with charge-altering releasable transporters elicits human T cell responses and cures established tumors in mice. Proc Natl Acad Sci USA 2018, 115 (39), E9153.

51. McKinlay, C. J.; Benner, N. L.; Haabeth, O. A.; Waymouth, R. M.; Wender, P. A., Enhanced mRNA delivery into lymphocytes enabled by lipid-varied libraries of charge-altering releasable transporters. Proc Natl Acad Sci U S A 2018, 115 (26), E5859–E5866.

52. Benner, N. L.; McClellan, R. L.; Turlington, C. R.; Haabeth, O. A. W.; Waymouth, R. M.; Wender, P. A., Oligo(serine ester) Charge-Altering Releasable Transporters: Organocatalytic Ring-Opening Polymerization and their Use for in Vitro and in Vivo mRNA Delivery. J. Am. Chem. Soc. 2019, 141 (21), 8416–8421.

53. Haabeth, O. A. W.; Blake, T. R.; McKinlay, C. J.; Tveita, A. A.; Sallets, A.; Waymouth, R. M.; Wender, P. A.; Levy, R., Local Delivery of Ox40l, Cd80, and Cd86 mRNA Kindles Global Anticancer Immunity. Cancer Res. 2019, 79 (7), 1624–1634.

54. Habibian, M.; McKinlay, C.; Blake, T. R.; Kietrys, A. M.; Waymouth, R. M.; Wender, P. A.; Kool, E. T., Reversible RNA acylation for control of CRISPR–Cas9 gene editing. Chemical Science 2020, 11 (4), 1011–1016.

55. Testa, S.; Haabeth, O. A. W.; Blake, T. R.; Del Castillo, T. J.; Czerwinski, D. K.; Rajapaksa, R.; Wender, P. A.; Waymouth, R. M.; Levy, R., Fingolimod-Conjugated Charge-Altering Releasable Transporters Efficiently and Specifically Deliver mRNA to Lymphocytes In Vivo and In Vitro. Biomacromolecules 2022, 23 (7), 2976–2988.

56. Wilk, A. J.; Weidenbacher, N. L.-B.; Vergara, R.; Haabeth, O. A. W.; Levy, R.; Waymouth, R. M.; Wender, P. A.; Blish, C. A., Charge-altering releasable transporters enable phenotypic manipulation of natural killer cells for cancer immunotherapy. Blood Advances 2020, 4 (17), 4244–4255.

57. Blake, T. R.; Haabeth, O. A. W.; Sallets, A.; McClellan, R. L.; Del Castillo, T. J.; Vilches-Moure, J. G.; Ho, W. C.; Wender, P. A.; Levy, R.; Waymouth, R. M., Lysine-Derived Charge-Altering Releasable Transporters: Targeted Delivery of mRNA and siRNA to the Lungs. Bioconjug Chem 2023, 34, 673–685.

58. Chen, R.; Wang, S. K.; Belk, J. A.; Amaya, L.; Li, Z.; Cardenas, A.; Abe, B. T.; Chen, C.-K.; Wender, P. A.; Chang, H. Y., Engineering circular RNA for enhanced protein production. Nature Biotechnology 2023, 41 (2), 262–272.

59. Blake, T. R.; Ho, W. C.; Turlington, C. R.; Zang, X. Y.; Huttner, M. A.; Wender, P. A.; Waymouth, R. M., Synthesis and mechanistic investigations of pH-responsive cationic poly(aminoester)s. Chemical Science 2020, 11 (11), 2951–2966.

60. Dilliard, S. A.; Cheng, Q.; Siegwart, D. J., On the mechanism of tissue-specific mRNA delivery by selective organ targeting nanoparticles. Proc Natl Acad Sci USA 2021, 118 (52), e2109256118.

61. Li, Z.; Amaya, L.; Ee, A.; Wang, S. K.; Ranjan, A.; Waymouth, R. M.; Chang, H. Y.; Wender, P. A., Organ– and Cell-Selective Delivery of mRNA In Vivo Using Guanidinylated Serinol Charge-Altering Releasable Transporters. J. Am. Chem. Soc. 2024, 146 (21), 14785–14798.

62. Ramírez Marrero, I. A.; Boudreau, L.; Hu, W.; Gutzler, R.; Kaiser, N.; von Vacano, B.; Konradi, R.; Perry, S. L., Decoupling the Effects of Charge Density and Hydrophobicity on the Phase Behavior and Viscoelasticity of Complex Coacervates. Macromolecules 2024, 57 (10), 4680–4694.

63. Hazra, B.; Mondal, A.; Prasad, M.; Gayen, S.; Mandal, R.; Sardar, A.; Tarafdar, P. K., Lipidated Lysine and Fatty Acids Assemble into Protocellular Membranes to Assist Regioselective Peptide Formation: Correlation to the Natural Selection of Lysine over Nonproteinogenic Lower Analogues. Langmuir 2022, 38 (49), 15422–15432.

64. Hazra, B.; Prasad, M.; Roy, R.; Tarafdar, P. K., The microenvironment and pK(a) perturbation of aminoacyl-tRNA guided the selection of cationic amino acids. Org Biomol Chem 2021, 19 (37), 8049–8056.

65. Blake, T. R.; Ho, W. C.; Turlington, C. R.; Zang, X.; Huttner, M. A.; Wender, P. A.; Waymouth, R. M., Synthesis and mechanistic investigations of pH-responsive cationic poly(aminoester)s. Chemical Science 2020, 11 (11), 2951–2966.

66. McKinlay, C. J.; Vargas, J. R.; Blake, T. R.; Hardy, J. W.; Kanada, M.; Contag, C. H.; Wender, P. A.; Waymouth, R. M., Charge-altering releasable transporters (CARTs) for the delivery and r elease of mRNA in living animals. Proceedings of the National Academy of Sciences 2017, 114 (4), E448–E456.

67. Li, S.; Tseng, W. C.; Stolz, D. B.; Wu, S. P.; Watkins, S. C.; Huang, L., Dynamic changes in the characteristics of cationic lipidic vectors after exposure to mouse serum: implications for intravenous lipofection. Gene Therapy 1999, 6 (4), 585–594.

68. Wright, M. J.; Rosenthal, E.; Stewart, L.; Wightman, L. M. L.; Miller, A. D.; Latchman, D. S.; Marber, M. S., β-Galactosidase staining following intracoronary infusion of cationic liposomes in the in vivo rabbit heart is produced by microinfarction rather than effective gene transfer: a cautionary tale. Gene Therapy 1998, 5 (3), 301–308.

69. Moghimi, S. M.; Symonds, P.; Murray, J. C.; Hunter, A. C.; Debska, G.; Szewczyk, A., A two-stage poly(ethylenimine)-mediated cytotoxicity: implications for gene transfer/therapy. Molecular Therapy 2005, 11 (6), 990–995.

70. Breunig, M.; Lungwitz, U.; Liebl, R.; Goepferich, A., Breaking up the correlation between efficacy and toxicity for nonviral gene delivery. Proc Natl Acad Sci USA 2007, 104 (36), 14454–14459.

71. Omo-Lamai, S.; Zamora, M. E.; Patel, M. N.; Wu, J.; Nong, J.; Wang, Z.; Peshkova, A.; Majumder, A.; Melamed, J. R.; Chase, L. S.; Essien, E. O.; Weissman, D.; Muzykantov, V. R.; Marcos-Contreras, O. A.; Myerson, J. W.; Brenner, J. S., Physicochemical Targeting of Lipid Nanoparticles to the Lungs Induces Clotting: Mechanisms and Solutions. Adv Mater 2024, 36 (26), e2312026.

72. Pert, E. K.; Hurst, P. J.; Waymouth, R. M.; Rotskoff, G. M., Coacervation drives morphological diversity of mRNA encapsulating nanoparticles. The Journal of Chemical Physics 2025, 162 (7).

73. Putnam, C. D.; Hammel, M.; Hura, G. L.; Tainer, J. A., X-ray solution scattering (SAXS) combined with crystallography and computation: defining accurate macromolecular structures, conformations and assemblies in solution. Quarterly Reviews of Biophysics 2007, 40 (3), 191–285.

74. Rosenberg, D. J.; Hura, G. L.; Hammel, M., Chapter Six – Size exclusion chromatography coupled small angle X-ray scattering with tandem multiangle light scattering at the SIBYLS beamline. In Methods Enzymol., Tainer, J. A., Ed. Academic Press: 2022; Vol. 677, pp 191–219.

75. Simonsen, J. B., A perspective on bleb and empty LNP structures. Journal of Controlled Release 2024, 373, 952–961.

76. Huertas, A.; Guignabert, C.; Barberà, J. A.; Bärtsch, P.; Bhattacharya, J.; Bhattacharya, S.; Bonsignore, M. R.; Dewachter, L.; Dinh-Xuan, A. T.; Dorfmüller, P.; Gladwin, M. T.; Humbert, M.; Kotsimbos, T.; Vassilakopoulos, T.; Sanchez, O.; Savale, L.; Testa, U.; Wilkins, M. R., Pulmonary vascular endothelium: the orchestra conductor in respiratory diseases. European Respiratory Journal 2018, 51 (4), 1700745.

77. Weber, E.; Sozio, F.; Borghini, A.; Sestini, P.; Renzoni, E., Pulmonary lymphatic vessel morphology: a review. Ann Anat 2018, 218, 110–117.

78. Townsley, M. I., Structure and composition of pulmonary arteries, capillaries, and veins. Compr Physiol 2012, 2 (1), 675–709.

79. Gillich, A.; Zhang, F.; Farmer, C. G.; Travaglini, K. J.; Tan, S. Y.; Gu, M.; Zhou, B.; Feinstein, J. A.; Krasnow, M. A.; Metzger, R. J., Capillary cell-type specialization in the alveolus. Nature 2020, 586 (7831), 785–789.

80. Madisen, L.; Zwingman, T. A.; Sunkin, S. M.; Oh, S. W.; Zariwala, H. A.; Gu, H.; Ng, L. L.; Palmiter, R. D.; Hawrylycz, M. J.; Jones, A. R.; Lein, E. S.; Zeng, H., A robust and high-throughput Cre reporting and characterization system for the whole mouse brain. Nat Neurosci 2010, 13 (1), 133–40.

81. Dirisala, A.; Uchida, S.; Li, J.; Van Guyse, J. F. R.; Hayashi, K.; Vummaleti, S. V. C.; Kaur, S.; Mochida, Y.; Fukushima, S.; Kataoka, K., Effective mRNA Protection by Poly(l-ornithine) Synergizes with Endosomal Escape Functionality of a Charge-Conversion Polymer toward Maximizing mRNA Introduction Efficiency. Macromolecular Rapid Communications 2022, 43 (12), 2100754.

