## Supplemental info for "*In vivo* mRNA delivery to the lung vascular endothelium by dicationic Charge-Altering Releasable Transporters"

[a] Department of Chemistry, Stanford University, Stanford, CA 94305, United States.

### Supporting Information

#### Materials and methods

**Materials:** Deuterated solvents ($D_2O$, $CD_3OD$, $CDCl_3$) were purchased from Cambridge Isotope Laboratories and used without further purification. Dialysis bags (regenerated cellulose, 3.5 kD or 1 kD MWCO) were purchased from Spectra/Por. Nδ-Boc-L-ornithine methyl ester hydrochloride was purchased from AA Blocks. H-Dab(boc) methyl ester hydrochloride and H-Dap(boc) methyl ester hydrochloride were purchased from Ambeed. Di-*tert*-butyl dicarbonate was purchased from AK scientific. *p*-Toluenesulfonic acid and (S)-(-)-3-amino-2-piperidone was purchased from TCI America. The MTC dodecyl monomer was prepared according to previously published methods.[1] All other substrates were purchased from Sigma Aldrich. fLuc-encoding mRNA (L-6107) and Cy5-labeled fLuc-encoding mRNA (L-7702) were purchased from Trilink. Cy5-labeled eGFP-encoding mRNA was purchased from APExBIO (R1011). NanoLuc-encoding mRNA was provided by VLP Therapeutics. Nano-Glo assay substrate and buffer were purchased from Promega. *In vivo*-jetPEI was purchased from Polyplus and was complexed with mRNA at N:P = 8 according to kit instructions.

**Instrumentation:** $^1H$ NMR and $^{13}C$ NMR spectra were recorded on either a Varian 400 or an Inova 600 MHz spectrometer. $^{13}C$ NMR was conducted with $^1H$ decoupling. In $CDCl_3$ and $CD_3OD$, the chemical shifts were referenced to δ 7.26 or 3.31. In $D_2O$, the HMPA internal standard was referenced to δ 2.61. Gel permeation chromatography (GPC) was performed in tetrahydrofuran (THF) at a flow rate of 1.0 ml/min on a Shimadzu chromatograph equipped with two columns (Jordi DVB 1000 Å and DVB Mixed Bed Low, temperature 40 $^oC$) connected in series. A Wyatt Optilab T-rEX refractive index detector and a Wyatt miniDAWN three-angle light scattering detector were employed. Cellular luminescence was measured using a SpectraMax L or SpectraMax iD3 microplate reader. Flow cytometry was performed using a Cytek Aurora or a NovoCyte Penteon (Stanford University Shared FACS Facility). FACS was performed using a BD FACSymphony S6 Sorter (Stanford University Shared FACS Facility).

**Polymerization:** Unless otherwise noted, polymerizations were carried out in glove boxes under $N_2$ atmosphere. For low temperature polymerizations, stock solutions and/or reaction mixtures were prepared in glove boxes, then removed and cooled to -78 $^oC$ with a dry ice/acetone bath over a stir plate, and quenched at low temperature with the specified quenching agent before being warmed to room temperature.

**Dynamic light scattering (DLS) and electrophoretic light scattering (ELS):** Dynamic Light Scattering (DLS) and Zeta Potential Measurements (Electrophoretic Light Scattering): DLS was measured on a Malvern Zetasizer. Following formulation, the CART/mRNA suspensions were diluted with 0.8 mL of RNase free ultrapure water and immediately measured in a disposable cuvette at an angle of 173°. Zeta potential was measured following DLS measurements by transferring the diluted CART/mRNA solutions into an electrophoretic cell and measured three times. Each sample was prepared three times and measured, and the average data is reported.
**Encapsulation efficiency:** Encapsulation efficiency (EE) was measured by using the RiboGreen assay (Invitrogen™ Quant-iT™ RiboGreen RNA Kit (ThermoFisher Scientific, Cat # R11490). Formulated CART-fLuc mRNA nanoparticles were prepared by addition and vortexing of CARTs (5 mM polymer in DMSO) to fLuc mRNA in PBS 5.5 (1.5 µg mRNA in 30 uL total

volume) with a 10:1 (+/-) ratio. Formulated solutions were diluted into the wells of the 96 well plate to reach a final concentration of 150 µg/mL with TE buffer. A total mRNA (mRNATotal) standard was prepared from a mRNA PBS solution (1.5 µg mRNA in 30 uL total volume) prepared without CART and diluted analogously to 150 µg/mL with TE buffer. The samples were incubated with 100 uL of 100-fold diluted Quant-iT™ RiboGreen™ RNA Reagent with gentle mixing for 5 minutes. The fluorescence from each sample was measured (excitation wavelength: 465 nm, Emission wavelength: 528 nm) on a SpectraMax iD3 Microplate Reader. The encapsulation efficiency was calculated as: $\boldsymbol{EE} = \frac{mRNA_{Free}}{mRNA_{Total}}$. Encapsulation efficiencies are reported as an average of two independently formulated CART/mRNA nanoparticle samples.

**Cryogenic transmission electron microscopy (cryoEM):** CryoEM samples were prepared from freshly prepared **CART**/mRNA suspensions onto Quantifoil R1.2/1.3 + 2 nm carbon (Electron Microscopy Sciences) grids. Grids were glow-discharged for 15 s at 10 mA to increase the hydrophilicity prior to sample loading. Vitrification was carried out by an Automatic Plunge Freeze Vitrobot Mark IV (Thermo Scientific) with 3 µL of the sample. Grid preparation was performed at 100% humidity, and the grids were blotted for 3 s at blot force 1 prior to plunging into liquid ethane. CryoEM samples were then imaged on a Glacios transmission electron microscope operating at 200 keV. Images were recorded using Serial EM software in low dose imaging mode at 45k magnification with a direct electron detector (K3).

**Small-angle X-ray scattering (SAXS):** SAXS measurements were carried out at the Advanced Light Source (ALS) SIBYLS (Structurally Integrated Biology for Life Sciences) beamline (12.3.1). Samples were prepared prior to measurement on site in 30 µL PBS (pH = 5.5) buffer-based formulations with a final RNA concentration of 1 mg/mL and a corresponding equivalence of CART to achieve N:P ratios ranging from 1:1 to 10:1. Samples were measured 30 times, in short segments with measurements taking approximately one minute to complete. All measurements were background subtracted from a PBS pH=5.5 sample. Runs 1 and 30 were compared to see if any beam damage or artefacts had occurred. No beam damage was detected in any of the samples. All runs were then averaged and plotted.

**Cell culture:** HeLa cells and A549 cells were maintained in Dulbecco's Modified Eagle Medium (DMEM) (Thermo Fisher Scientific; high glucose, pyruvate) supplemented with 10% (v/v) fetal bovine serum (FBS) and 1% penicillin/streptomycin. Cells were passaged at ~80% confluence.

**Cy5-labeled eGFP mRNA delivery into A549 cells:** A549 cells were seeded at 60,000 cells/well in 800 µl DMEM containing 10% FBS and 1% penicillin/streptomycin in a 24-well plate. Cells were incubated 22 h at 37 ˚C (5% $CO_2$) and then washed with serum-free DMEM, and then to each well was added 300 µl serum-free DMEM. mRNA/CART particles were prepared at a 10:1 (+/-) charge ratio. Briefly, to 16.15 µl pH 5.5 PBS in a 600 µl microcentrifuge tube was added 6.0 µl Cy5-labeled eGFP-encoding mRNA (stored as a 0.2 µg/µl solution in pH 7.4 PBS). To this was added 1.85 µl D-**Orn-CART** (D16:Orn5, stored as a 2.0 mM solution in DMSO). This was mixed for 20 s, at which point 3 µl of solution was added to each of three wells (resulting in a final dose of 150 ng mRNA/well). The cells were incubated at 37 ˚C for 16.5 h, at which point the media was removed and the cells were trypsinized with 0.25% trypsin-EDTA and transferred to a 96-well round-bottom plate. The plate was then centrifuged, the supernatant was removed, and the pellets were re-dispersed in FACS buffer (100 µl/well). Centrifugation, supernatant removal, and resuspension in FACS buffer was repeated two additional times, and then the plate was submitted for flow cytometry (NovoCyte Penteon, Stanford University Shared FACS Facility). Singlet cell populations for each treatment were analyzed for eGFP and Cy5 median fluorescence intensity. Data are presented as representative examples or as the average of 2-3 wells. Error is expressed as ± one standard deviation.

***In vivo* Fluc mRNA delivery studies:** All mice used in in vivo studies were Female BALB/c mice housed in the Laboratory Animal Facility of the Stanford University Medical Center. Experimental protocols were approved by the Stanford Administrative Panel on Laboratory Animal Care. **CART**/fLuc mRNA particles were formulated as described above using 5 µg mRNA in a total formulation volume of 100 µl per mouse. For example, for D-**Orn-CART** at a 10:1 (+/-) charge ratio: to 110.3 µl pH 5.5 PBS in a 1.5 ml microcentrifuge tube was added 6.0 µl fLuc mRNA (stored as a 1.0 µg/µl solution). To this was added 3.7 µl D-**Orn-CART** (D15:Orn5, stored as a 5.0 mM solution in DMSO). This was mixed for 20 s, at which point 100 µl of the 120 µl solution (dose of 5 µg mRNA) was injected intravenously via the tail vein. After 4 h, mice were anesthetized with isoflurane and 150 mg/kg D-luciferin was injected intraperitoneally. After waiting 5-7 minutes, luminescence was measured using an AMI imaging system. For imaging of excised organs, mice were sacrificed after full-body imaging. Lungs, spleen, liver, and kidneys of each mouse were collected and imaged within 5 minutes of sacrifice. For imaging of excised organs of mice treated with NanoLuc mRNA, lungs, spleen, and liver of each mouse were collected and incubated with a solution of D-luciferin for 5 minutes, then removed from the solution, blotted dry, and imaged.

**Cre mRNA delivery into reporter mice to identify CART-targeted cell types *in situ* :** Cre reporter mice containing a ubiquitously driven *loxP*-flanked STOP cassette followed by tdTomato cDNA were obtained from the Jackson Laboratory (strain 007914) and crossed onto a BALB/c background from Charles River (strain 028) for seven generations before heterozygotes were crossed to generate a homozygous breeding stock. Cre mRNA was obtained from Trilink Biotechnologies and delivered using D-Orn-CART and D-Daba-CART by tail vein injection in a total of 3 mice per CART, 5µg per animal. One un-injected littermate was retained in each experiment to control for Cre-independent tdTomato expression.

**Tissue Collection**: Animals were euthanized by $CO_2$ inhalation 2 days after CART injections. Blood was washed from the pulmonary circulation by slow (<100ml per 10 seconds) perfusion of PBS into the right ventricle of the heart. Airways were inflated with 2% low melting point agarose (Invitrogen, 16520) prepared in sterile PBS. Kidneys, liver, spleen, heart and lungs were harvested and submerged in 40ml ice-cold 4% paraformaldehyde solution in PBS (EMS, 15710) for 3hrs at 4°C with gentle shaking protected from light, after which tissue was washed briefly in PBS.

**Histology**: Vibratome sections were cut immediately to a thickness of 300mm using a Leica VT1000 S vibratome and were stored in PBS at 4°C protected from light. Cryosections were prepared following standard protocols. Briefly, 4% PFA fixed lobes were cryopreserved in 30%

sucrose in PBS, embedded in OCT (Sakura, 4583) and 25mm sections were cut using a Leica CM3050S cryostat. Cut sections were stored at -80°C prior to staining.

**Immunohistochemistry on cryosections**: Cryosections were stained following standard IHC protocols as described in previous publications[3]. Briefly, slides were thawed, washed in PBS + 0.1% Tween-20, blocked at least 30 min in preblock consisting of 0.3% Triton X-100, 5% serum and 1.5% BSA, and then incubated overnight in primary antibody solution diluted in preblock at room temperature. The next day slides were washed, incubated for 45 minutes in secondary antibody solution, nuclei were stained with DAPI (1:1000 dilution of 10mg/ml stock; Invitrogen, D1306), and slides were mounted with Prolong Gold. For visualization of elastin fibers by staining with fluorescent hydrazide dye, stock solutions were prepared by dissolving 1mg hydrazide-A633 (Invitrogen, A30634) in 2ml di$H_2O$, and were used at a 1:500 (A633) dilution in the primary antibody mix. Ordering and dilution information for the primary antibodies used: mouse anti SMA-FITC to identify smooth muscle a-actin expressing cells (Sigma, F3777; 1:200), hamster anti mouse CD31 (BioRad, MCA1370Z; 1:200), Rabbit anti ERG to identify endothelial nuclei (Abcam, ab110639; 1:100).

**Antibody staining and tissue clearing of vibratome sections**: Vibratome sections were stained as described previously. Briefly, vibratome sections floating freely in a round bottom 5ml tube were incubated first in preblock solution as above, then in primary antibody solution diluted in preblock for 1-3 nights at 4 degrees with gentle shaking, protected from light. Sections were then washed briefly in PBS with 0.1% Tween-20 and stored at 4 degrees protected from light until ready for imaging. Individual vibratome sections were cleared in Cubic 1 and mounted on glass slides immediately prior to confocal imaging.

**Confocal imaging, image analysis and statistical analysis**: All confocal images were captured on a Zeiss 880 Examiner confocal microscope and maximum intensity projections were made and minimally processed using Zen Black software (Carl Zeiss AG). Tiled z stacks were used to capture large regions or structures at high resolution. Measurements of the tdTomato-marked fraction of lung artery and vein endothelial cells and percentage of tdTomato-marked ERG+ capillary nuclei were performed manually with the assistance of Zen Black. Statistical tests used are detailed in figure legends.

**Blood chemistry analysis:** mice were either left untreated or treated as described above with Orn-CART formulated with 5 µg fLuc mRNA at a 10:1 (+/-) charge ratio. For acute toxicity study, blood was collected 24 h after treatment, mice were anesthetized and blood was collected in EDTA-containing tubes. For chronic toxicity study, mice were injected weekly for 4 weeks (total 4 doses) and blood was collected 24 hr after last dose. Chemistry analysis was performed at the Stanford Animal Diagnostic Laboratory on the Siemens Dimension EXL200/LOCI analyzer. A clinical laboratory scientist performed all testing, including dilutions and repeat tests as indicated, and reviewed all data.

**Measurement of NanoLuc protein expression in lung cell populations: Orn-CART**: NanoLuc mRNA particles were formulated at a 10:1 (+/-) charge ratio as described above using 10 µg NanoLuc mRNA in a total formulation volume of 100 µl per mouse. For example: to 98.5

µl pH 5.5 PBS in a 1.5 ml microcentrifuge tube was added 14.2 µl NanoLuc-encoding mRNA (stored as a 0.848 µg/µl solution). To this was added 7.4 µl **Orn-CART** (D15:Orn5, stored as a 5.0 mM solution in DMSO). This was mixed for 20 s, at which point 100 µl of the 120 µl solution (dose of 10 µg mRNA) was injected intravenously via the tail vein. After 16 h, mice were sacrificed, perfused with 10 mL cold PBS, and lungs were collected. Lungs were minced with scissors and digested in a solution of collagenase D/hyaluronidase (STEMCELL Technologies kit) for 30 minutes at 37 °C, then passed through a 70 um cell strainer to yield single cell suspensions. Cell suspensions were incubated in ACK lysis buffer for 5 minutes and then washed and stained using Live/Dead Fixable Blue Stain, FITC-CD31, BUV379-CD45, and BV421-CD326. Cells were sorted into epithelial cell (CD45- epCAM+), endothelial cell (CD45- CD31+), and immune cell (CD45+) populations (10k cells/population) using a BD FACSymphony S6 Sorter (Stanford University Shared FACS Facility). Each population was transferred to one well of a black-walled 96-well plate and 50 µl Nano-Glo substrate solution (1:50 v:v substrate:lysis buffer) was added to each well. After 5 minutes, the cells were read on a SpectraMax L microplate reader. Data are presented as the average of 3 wells. Error is expressed as ± one standard deviation.

**Statistical analysis:** Statistical analysis between groups was performed using an unpaired student t-test (GraphPad Prism 9). Statistical tests used for each figure are detailed in figure legends.

### Supporting Information

#### Synthetic procedures

**Synthesis of $M_{Orn}$ precursor methyl 2-((*tert*-butoxycarbonyl)(2-hydroxyethyl)amino)-5-((*tert*-butoxycarbonyl)amino)pentanoate**

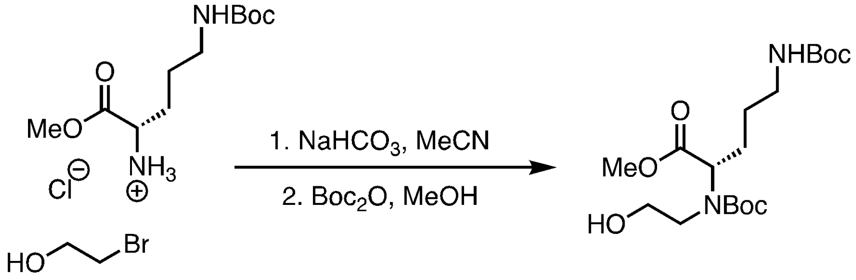


A solution of Nδ-Boc-L-ornithine methyl ester hydrochloride (2.0 g, 7.07 mmol, 1.0 eq) and sodium bicarbonate (1.54 g, 18.38 mmol, 2.6 eq) in acetonitrile (21 ml) in a round-bottom flask was refluxed with stirring for 20 minutes. 2-Bromoethanol (601 µl, 8.49 mmol, 1.2 eq) was added in one shot. The reaction was stirred at reflux for 22 hours, until the consumption of 2-bromoethanol plateaued, as determined by $^{1}H$ NMR. The crude product was cooled, diluted in acetone, and filtered through celite. The solution was concentrated to dryness under reduced pressure and the crude product was resuspended in methanol (7.8 ml). The solution was sparged with nitrogen for 5 minutes, and then to the solution was added di-tert-butyl dicarbonate (1.62 ml, 7.07 mmol, 1.0 eq). The reaction was stirred at room temperature for 19 hours. The solution was concentrated to dryness under reduced pressure to yield the crude product as a yellow oil. Purification by column chromatography (3:1 dichloromethane:ethyl acetate) yielded the product as a colorless oil (1.60 g, 58% yield). Purity confirmed by $^{1}H$ NMR (Figure S18, S19).

$^{1}H$ NMR ($CDCl_3$, 400 MHz): δ 4.75-4.47 (br, 1 H), 4.24-3.98 (m, 1H), 3.89-3.46 (m, 6 H), 3.27-3.07 (m, 3 H), 2.13-1.82 (m, 2H), 1.72-1.38 (m, 21 H)

$^{13}C$ NMR ($CDCl_3$, 600 MHz): δ 173.75, 173.06, 156.09, 155.47, 81.19, 80.83, 79.26, 61.78, 61.49, 61.35, 60.62, 52.60, 52.49, 51.32, 50.47, 40.15, 28.45, 28.34, 27.48, 27.13, 26.55.

**Synthesis of ornithine-derived morpholinone ($M_{Orn}$) *tert*-butyl 3-(3-((*tert*-butoxycarbonyl)amino)propyl)-2-oxomorpholine-4-carboxylate**

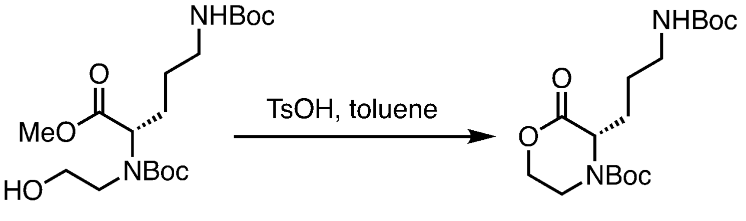


**$M_{Orn}$**

A toluene solution of the isolated di-Boc-protected N-hydroxyl ornithine methyl ester (1.59 g, 4.08 mmol) and *p*-toluenesulfonic acid monohydrate (101 mg, 0.53 mmol, 0.13 eq) was heated to reflux. The reaction was stirred at reflux for 45 minutes, after which the solution was concentrated to dryness. Purification by column chromatography (3:1 hexanes:ethyl acetate) yielded **$M_{Orn}$** (1.16 g, 79% yield) as a viscous, white oil. Purity confirmed by $^1$H NMR (Figure S20, S21).

$^1$H NMR ($CDCl_3$, 400 MHz, Figure S20): δ 4.65 (br, 2H), 4.39 (t, 2H), 3.91 (br, 1H), 3.42 (br, 1H), 3.27-3.06 (m, 2H), 1.90 (m, 2H), 1.64 (m, 2H), 1.48 (s, 9H), 1.44 (s, 9H)

$^{13}$C NMR ($CDCl_3$, 400 MHz, Figure S21): δ 168.82, 155.08, 153.81, 81.62, 79.35, 67.48, 61.18, 39.99, 30.85, 28.50, 28.40, 27.06, 26.60

**Synthesis of $M_{Daba}$ precursor**

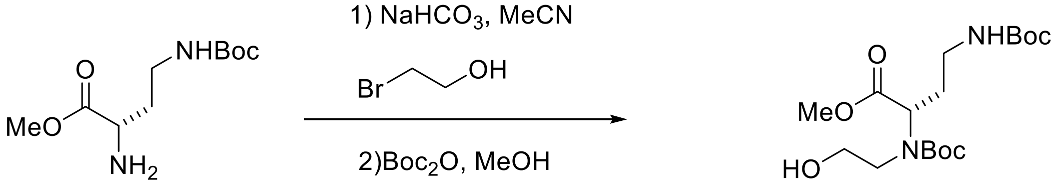


A solution of H-Dab(boc) methyl ester hydrochloride (Alternate name:(R)-Methyl 2-amino-4-((tert-butoxycarbonyl)amino)butanoate hydrochloride) (2.0 g, 7.44 mmol, 1.0 eq) and sodium bicarbonate (1.63 g, 19.35 mmol, 2.6 eq) in acetonitrile (21 ml) in a round-bottom flask was refluxed with stirring for 20 minutes. 2-Bromoethanol (632 µl, 8.93 mmol, 1.2 eq) was added in one shot. The reaction was stirred at reflux for 22 hours, until the consumption of 2-bromoethanol plateaued, as determined by $^1$H NMR. The crude product was cooled, diluted in acetone, and filtered through celite. The solution was concentrated to dryness under reduced pressure and the crude product was resuspended in methanol (8.2 ml). The solution was sparged with nitrogen for 5 minutes, and then to the solution was added di-tert-butyl dicarbonate (1.71 ml, 7.44 mmol, 1.0 eq). The reaction was stirred at room temperature for 19 hours. The solution was concentrated to dryness under reduced pressure to yield the crude product as yellow oil. Purification by column chromatography (3:1 dichloromethane: ethyl acetate) yielded the product as a colorless oil (1.30 g, 46% yield).

$^1$H NMR (600 MHz, cdcl$_3$) δ 4.66 (s, 1H), 4.43 – 3.94 (m, 1H), 3.69 (br, 6H), 3.13 (m, 3H), 2.27 – 1.79 (m, 2H), 1.58 (s, 2H), 1.46 – 0.95 (m, 21H).
$^{13}$C NMR (101 MHz, $CDCl_3$) δ 173.29, 172.77, 156.08, 155.27, 81.21, 80.85, 79.38, 61.56, 61.37, 59.32, 58.54, 52.57, 52.47, 51.47, 50.72, 37.52, 31.02, 29.70, 28.35, 28.22.

##### Synthesis of Daba-derived morpholinone $M_{Daba}$

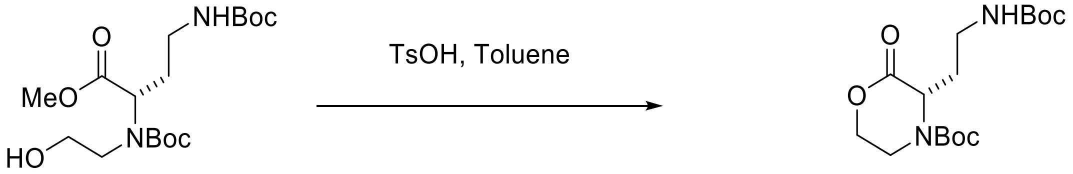


A solution of the isolated di-Boc-protected H-Dab methyl ester (1.30 g, 3.45 mmol) and *p*-toluenesulfonic acid monohydrate (85 mg, 0.45 mmol, 0.13 eq) was heated to reflux in toluene. The reaction was stirred at reflux for 45 minutes, after which the solution was concentrated to dryness. Purification by column chromatography (3:1 hexanes:ethyl acetate) yielded $\mathbf{M_{DABA}}$ (0.95 g, 80% yield) as a viscous, white oil.

$^1$H NMR (600 MHz, cdcl$_3$) δ 4.70 (s, 2H), 4.32 (t, *J* = 5.2 Hz, 2H), 3.81 (s, 1H), 3.12 (m, 3H), 2.15 – 1.73 (m, 2H), 1.40 (m, 18H).
$^{13}$C NMR (101 MHz, CDCl$_3$) δ 168.65, 155.74, 81.67, 79.16, 67.24, 53.69, 39.45, 36.71, 33.03, 28.33, 28.17.

##### Synthesis of DABA-derived homopolymer:

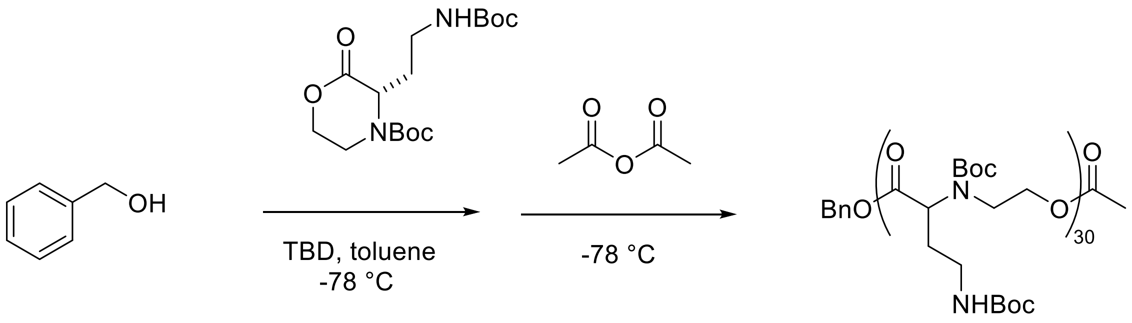


**p(HE-DABA)$_{Boc}$**

To a solution of DABA monomer $M_{DABA}$ (68.7 mg, 0.20 mmol, 30 eq) in toluene (150 μl) was added a solution of benzyl alcohol (0.69 μl, 0.0067 mmol, 1 eq) and 1,5,7-triazabicyclo[4.4.0]dec-5-ene (1.39 mg, 0.010 mmol, 5 mol% w/r/t monomer, 1.5 eq) in toluene (50 μL) under $N_2$. The reaction was stirred at room temperature for 45 seconds, after which it was cooled to -78 $^o$C and stirred for 20 minutes. A solution of acetic anhydride (31 μl, 0.33 mmol, 50 eq) in toluene (60 μl) under $N_2$ was added. The reaction was stirred for two minutes at -78 $^o$C and then warmed to room temperature and stirred to an additional 1 minute. The reaction mixture was concentrated to dryness. The crude mixture was resuspended in 2 ml of dichloromethane and dialyzed (RC tubing 1 kD MWCO) against methanol (800 ml). After 20 hours, p(HE-DABA)$_{Boc}$ was isolated as a white solid (19.6 mg, 28% yield). $^1$H NMR analysis indicates a degree of polymerization (DP) of 30.

GPC: dn/dc=0.076, $M_n$=5.7x10$^3$, Đ=1.06

$^1$H NMR (600 MHz, cdcl$_3$) δ 7.27 (s, 5H), 5.06 (s, 27H), 4.27 (m, 88H), 3.81 – 3.32 (m, 50H), 3.08 (m, 80H), 2.13 (m, 32H), 1.88 (s, 27H), 1.37 (m, 644H).

##### Synthesis of $M_{Dapa}$ precursor

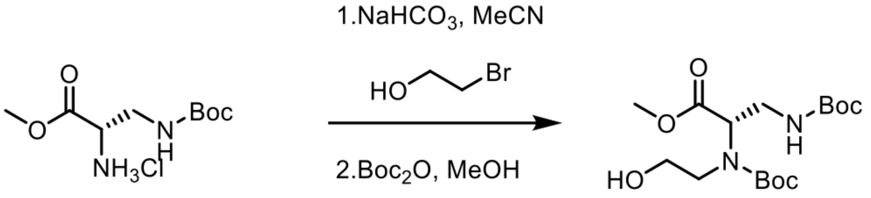


In round bottom flask equipped with a stir bar, H-Dap (Boc)-methyl ester hydrochloride (5.00g, 17.68 mmol, 1.00 eq), NaHCO3 (3.86g, 45.97 mmol, 2.60 eq), were dissolved in 500 mL MeCN and refluxed in an oil bath set to 50 degrees Celsius for 20 minutes. 2-Bromoethanol (2.65g, 21.22 mmol, 1.2 eq) diluted in 15 mL of MeCN was added over the course of one minute. The reaction was left to stir overnight with a reflux condenser. The crude product was cooled and filtered through celite with the aid of acetone. The solution was concentrated to dryness under reduced pressure and the crude product was resuspended in methanol (40 ml). The solution was sparged with nitrogen for 20 minutes, and then to the solution was added di-tert-butyl dicarbonate (4.06 mL, 17.68 mmol, 1.0 eq). The reaction was stirred at room temperature for 24 hours under nitrogen. The solution was concentrated to dryness under reduced pressure to yield the crude product as a yellow oil. Purification by column chromatography (10-50% ethyl acetate in dichloromethane) yielded roduct as a colorless oil (2.737 g, 43% yield). Purity confirmed by $^1$H NMR (Figure S18, S19). It was found that the $M_{Dapa}$ precursor degraded over time, and so it was taken forward in the synthesis of $M_{dapa}$ shortly after column chromatography and solvent removal

$^1$H NMR (600 MHz, cdcl$_3$) δ 5.07 (s, 1H), 4.11 – 3.98 (m, 1H), 3.92 – 3.84 (m, 1H), 3.83 – 3.67 (m, 5H), 3.67 – 3.23 (m, 3H), 1.51 – 1.39 (m, 18H).
$^{13}$C NMR (101 MHz, cdcl$_3$) δ 171.68, 171.43, 156.07, 155.95, 155.17, 154.69, 81.04, 80.64, 79.41, 61.11, 60.97, 52.25, 52.16, 51.65, 40.80, 40.03, 30.68, 28.16, 28.08, 28.05, 28.01.

##### Synthesis of Dapa-derived morpholinone $M_{Dapa}$

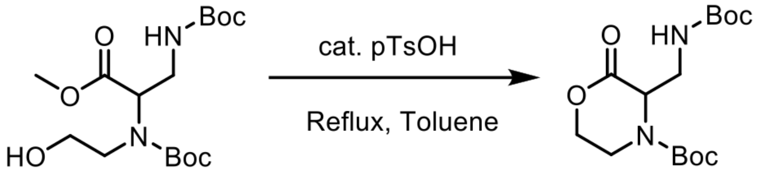


To a 500 mL round bottom flask charged with a magnetic stir bar was added toluene (80 mL) and p-toluenesulfonic acid monohydreate (0.187g, 0.98 mmol, 0.13 eq.). The solution was brought to a reflux. $M_{Dapa}$ precursor (2.737 g, 7.55 mmol, 1 eq.) was taken up in toluene (20 mL) and added to the refluxing solvent. A reflux condenser was attached and the reaction proceeded for 1 hour 30 minutes. The crude mixture was cooled at room temperature. Purification by column chromatography (15% ethyl acetate in dichloromethane) yielded the product after one more cycle as a colorless oil (1.242 g, 50% yield). Purity confirmed by 1H NMR.

$^{1}$H NMR (600 MHz, cdcl3) δ 5.08 – 4.75 (m, 1H), 4.75 – 4.50 (m, 1H), 4.49 – 4.26 (m, 2H), 4.05 – 3.67 (m, 1H), 3.67 – 3.47 (m, 2H), 3.39 (s, 1H), 1.60 – 1.24 (m, 18H).

$^{13}$C NMR (101 MHz, $CDCl_3$) δ 167.43, 155.82, 153.38, 81.51, 79.64, 67.93, 56.76, 55.65, 41.97, 39.86, 38.42, 28.20.

##### Synthesis of ornithine-derived copolymer Orn-CART$_{Boc}$

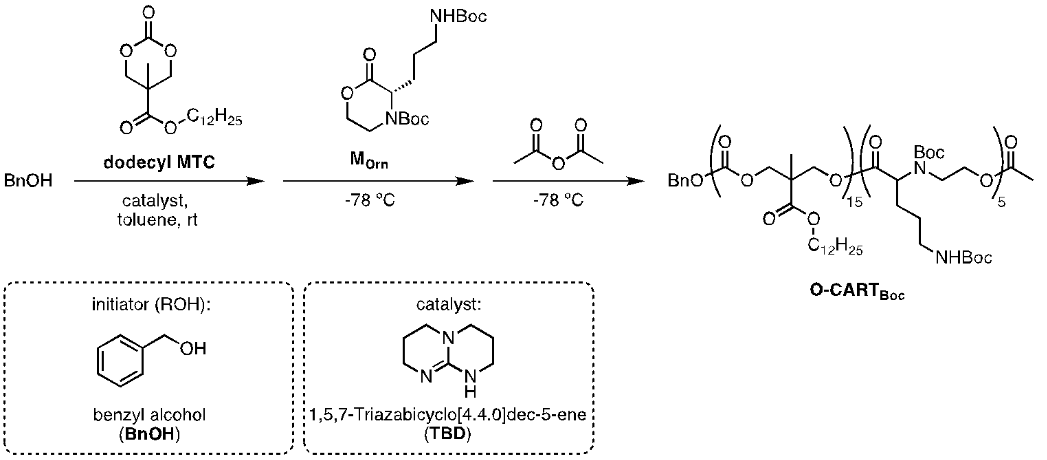


To a solution of **dodecyl MTC** (63.2 mg, 0.19 mmol, 13.75 eq) in toluene (70 µl) was added a solution of benzyl alcohol (1.44 µl, 0.014 mmol, 1 eq) and 1,5,7-triazabicyclo[4.4.0]dec-5-ene (1.34 mg, 0.0098 mmol, 0.7 eq) in toluene (20 µl) in a glovebox. This solution was stirred at room temperature. After 7 minutes, the solution of **dodecyl MTC**, initiator, and catalyst was added to **M$_{Orn}$** (25.0 mg, 0.07 mmol, 5.0 eq). The solution was cooled to -78 °C and stirred for 15 minutes, after which point a solution of acetic anhydride (66 µl, 0.7 mmol, 50 eq) in toluene (80 µl) was added. This was allowed to warm to room temperature. The crude residue was redissolved in dichloromethane and then dialyzed (regenerated cellulose tubing, MWCO 1kD) against methanol for 18 hours to yield **Orn-CART$_{Boc}$** as a clear oil (69.9 mg). $^{1}$H NMR analysis (CDCl3, 400 MHz) reveals an oligomer with 15 dodecyl units and 5 ornithine units. These DPs were determined by comparing the aromatic initiator signal (7.39-7.30 ppm, 5H) to the signal of the **M$_{Orn}$** block (3.21-3.01 ppm, 10 H), and signal from dodecyl block (0.93-0.80 ppm, 44 H). Note: in this manuscript, we define "**Orn-CART**" as the oligomer with m = 15 or 16, n = 5; that is, BnO-$D_{15}$-*b*-$Orn_5$-OAc or BnO-$D_{16}$-*b*-$Orn_5$-OAc.
GPC: dn/dc=0.042, $M_n$=5.6x10$^3$, Đ=1.68
$^{1}$H NMR ($CDCl_3$, 400 MHz, Figure S30): δ 7.39-7.30 (m, 5 H), 5.18-5.10 (s, 2H),iii (br, 4H), 4.48-4.16 (m, 67H), 4.16-4.02 (m, 33H), 3.81 (m, 1.5H), 3.64-3.21 (m, 10H), 3.21-3.01 (br, 10H), 2.08-1.93 (m, 10H), 1.90-1.71 (br, 11 H), 1.68-1.57 (t, 30H), 1.58-1.50 (br, 10H), 1.50-1.37 (m, 85H), 1.37-1.09 (m, 308H), 0.93-0.80 (t, 44H)

#### Deprotection of Orn-CART$_{Boc}$ to form Orn-CART

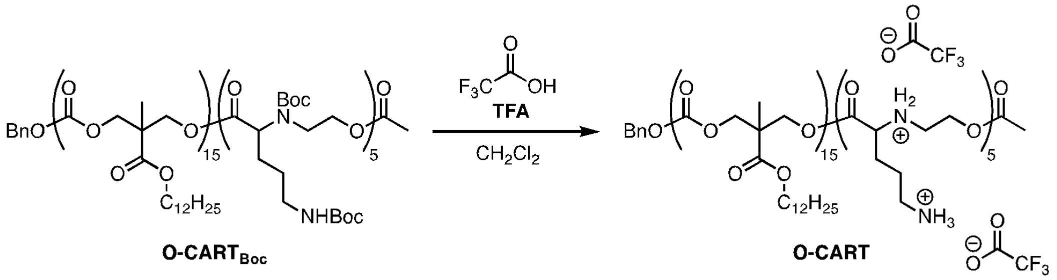


To a solution of **Orn-CART$_{Boc}$** (n = 15, m = 5, 10.8 mg) in dry, degassed dichloromethane (1.08 ml) was added trifluoroacetic acid (108 µl). The solution was stirred under a nitrogen atmosphere for 4 hours, after which point the product was concentrated to dryness to yield a clear oil (12.1 mg). The product **Orn-CART** was analyzed by $^1$H NMR ($CD_3OD$, 600 MHz) and then dissolved in dimethyl sulfoxide to make a 5 mM solution, which was used in *in vitro* and *in vivo* experiments without further purification.

$^1$H NMR ($CD_3OD$, 600 MHz): δ 7.42-7.26 (m, 5H), 5.34-5.28 (s, 1H), 5.17-5.06 (s, 2H), 4.49-4.20 (m, 66H), 4.20-4.02 (m, 40H), 3.45-3.30 (m, 9H), 3.05-2.87 (m, 9H), 2.09-1.96 (m, 14H), 1.95-1.68 (m, 12H), 1.69-1.51 (s, 32H), 1.46-1.07 (m, 340H), 0.93-0.83 (t, 48H).

#### Synthesis of Oleyl Ornithine-derived copolymer Oleyl-Orn-CART$_{Boc}$

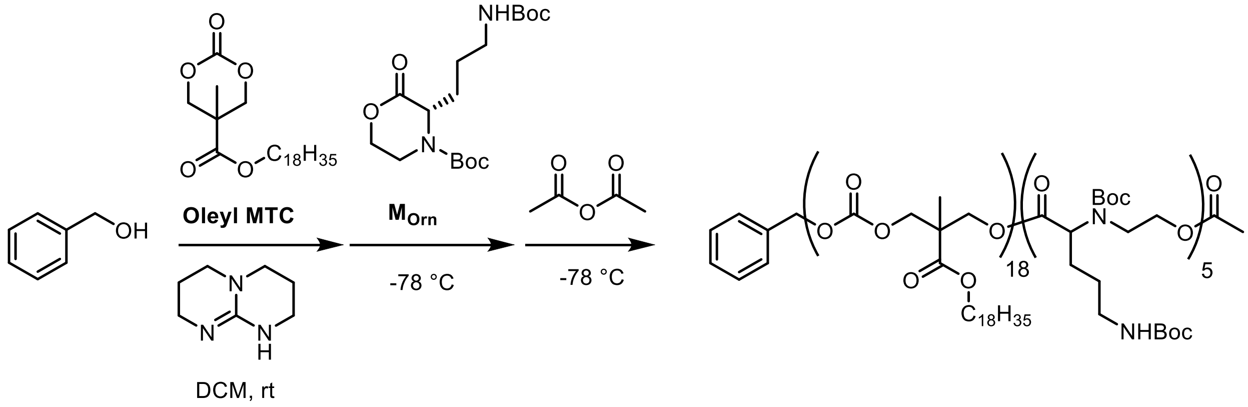


To a solution of **Oleyl MTC** (264.8 mg, 0.645 mmol, 13.75 eq) in dichloromethane (240 µl) was added a solution of benzyl alcohol (5.07 mg, 0.047 mmol, 1 eq) and 1,5,7-triazabicyclo[4.4.0]dec-5-ene (4.57 mg, 0.033 mmol, 0.7 eq) in dichloromethane (60 µl) in a glovebox. This solution was stirred at room temperature. After 7 minutes, the reaction solution was added to **M$_{Orn}$** (68.3 mg, 0.191 mmol, 4.06 eq). The solution was cooled to -78 $^o$C and stirred for 15 minutes, after which point a solution of acetic anhydride (0.1 mL, x.s.). This was allowed to warm to room temperature. The crude residue was dialyzed (regenerated cellulose tubing, MWCO 1kD) against methanol overnight to yield **Oleyl-Orn-CART$_{Boc}$** as a clear oil (283.6 mg). $^1$H NMR analysis ($CDCl_3$, 400 MHz) reveals an oligomer with 18 Oleyl units and 5 Ornithine units. These DPs were determined by comparing the aromatic initiator signal (7.39-7.29 ppm, 5H) to the signal of the **M$_{Orn}$** block (3.20-3.05 ppm, 9 H), and signal from Oleyl block (0.87 ppm, 54H). Note: in this manuscript, we define "**Oleyl-Orn-CART**" as the oligomer with m = 18, n = 5; that is, BnO-$O_{15}$-*b*-$Orn_5$-OAc.
GPC: dn/dc=0.165, $M_n$=2.5 kDa, Đ=1.47

$^1$H NMR (400 MHz, cdcl$_3$) δ 7.36 (d, *J* = 3.4 Hz, 5H), 5.41 – 5.27 (m, 37H), 4.28 (s, 82H), 4.10 (q, *J* = 6.4 Hz, 43H), 3.70 (s, 7H), 3.13 (s, 9H), 2.01 (q, *J* = 6.4 Hz, 81H), 1.69 – 1.49 (m, 57H), 1.44 (d, *J* = 11.0 Hz, 86H), 1.29 (qd, *J* = 12.9, 5.6 Hz, 472H), 0.88 (t, *J* = 6.8 Hz, 54H).

#### Supporting Information

##### Rearrangement of p(HE-Orn)2+ at pH 6.5:

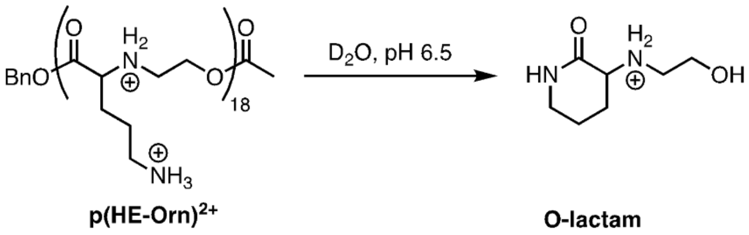


Deprotected polymer p(HE-Orn)$^{2+}_{18}$ (15.9 mg, 0.042 mmol by repeat units) was dissolved in 500 µL $D_2O$ with 3 µL HMPA (internal standard). The $t_0$ $^1H$ NMR spectrum was acquired with the parameters (Varian) nt = 8, at = 8, and d1 = 12, which were confirmed to provide a long enough time between scans to ensure accurate relative integrations. 500 µL of the 3.3 M ($HK_2PO_4/H_2KPO_4$) pH 6.5 buffer solution in $D_2O$ was then added to the NMR tube, which was inverted five times to ensure thorough mixing. NMR spectra were then acquired once every 90 s for 1.5 h using the parameters nt = 4, at = 8, and d1 = 12. The arrayed spectral data were analyzed using MNova software. The first time point was acquired at 170 seconds after buffer addition, which showed no conversion to lactam. The HMPA peak was referenced to 2.61 ppm. The concentration of (HE-Orn)$^{2+}$ repeat units was determined by relative integration of the peak at 3.15 ppm (integration 3.18-3.09, 2) relative to the HMPA standard. The formation of the ornithine lactam was monitored by formation of the peak at 4.06 ppm (integration 4.08-4.04 ppm, 1H). The p(HE-Orn)$^{2+}$ and O-lactam peaks at each timepoint were plotted relative to the p(HE-Orn)$^{2+}$ integration at 170 seconds to determine percent loss and percent conversion, respectively.

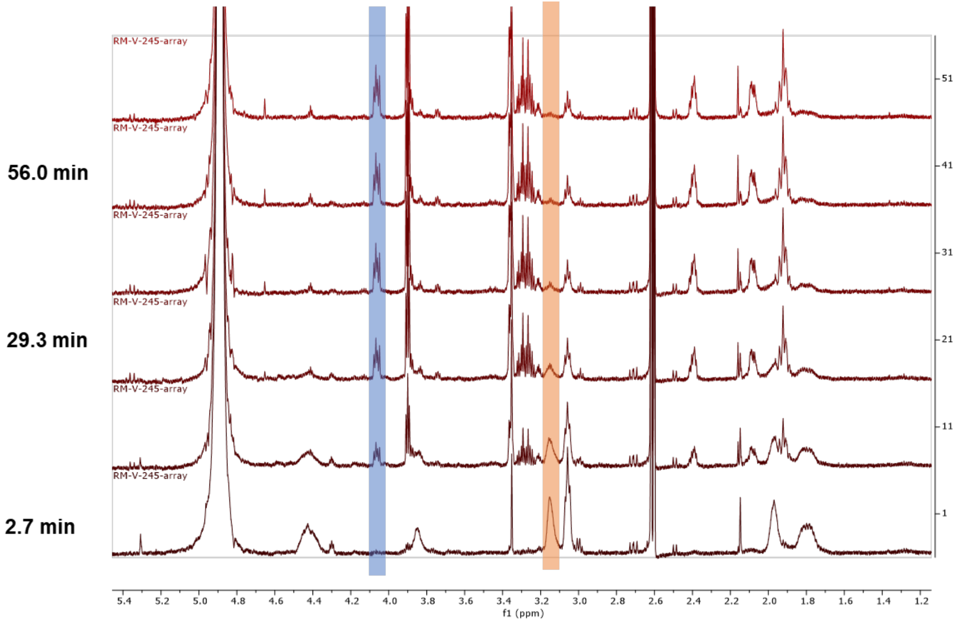


**Figure S1. Sample 1H NMR spectra of p(HE-Orn)2+ rearrangement at pH 6.5.** The peak highlighted in blue is the O-lactam and the peak highlighted in orange is the p(HE-Orn)$^{2+}$ repeat unit.

**Synthesis of Daba-derived copolymer (Daba-CART):**

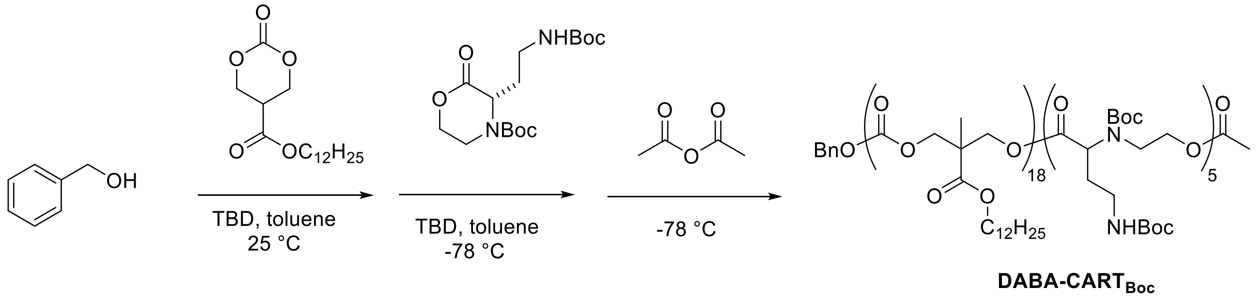


To a solution of **dodecyl MTC** (61.9 mg, 0.19 mmol, 11.1 eq) in toluene (85 µl) was added a solution of benzyl alcohol (1.75 µl, 0.017 mmol, 1 eq) and 1,5,7-triazabicyclo[4.4.0]dec-5-ene (1.66 mg, 0.0119 mmol, 0.7 eq) in toluene (20 µl) under a nitrogen atmosphere. This solution was stirred at room temperature. After 7 minutes, the solution of **dodecyl MTC**, initiator, and catalyst was added to $\mathbf{M_{Daba}}$ (29.2 mg, 0.085 mmol, 5.0 eq). The solution was cooled to -78 $^{o}$C and stirred for 15 minutes, after which point a solution of acetic anhydride (66 µl, 0.7 mmol, 50 eq) in toluene (80 µl) was added. This was allowed to warm to room temperature. The crude residue was redissolved in dichloromethane and then dialyzed (regenerated cellulose tubing, MWCO 1kD) against methanol for 18 hours to yield $\mathbf{Daba\text{-}CART_{Boc}}$ as a clear oil (35.3 mg). $^{1}$H NMR analysis (CDCl3, 400 MHz) reveals an oligomer with 18 dodecyl units and 5 DABA units. These DPs were determined by comparing the aromatic initiator signal (7.39-7.30 ppm, 5H) to the signal of the $\mathbf{M_{Daba}}$ block (3.51 ppm, 10 H), and signal from dodecyl block (0.93-0.80 ppm, 55 H). Note: in this manuscript, we define "**Daba -CART**" as the oligomer with m = 18, n = 5; that is, BnO-$D_{18}$-*b*-$Daba_5$-OAc.

GPC: dn/dc=0.042, $M_n$=5.6x$10^3$, Đ=1.68
$^{1}$H NMR (600 MHz, $cdcl_3$) δ 7.36 (s, 4H), 5.15 (s, 2H), 4.28 (s, 89H), 4.11 (s, 42H), 3.51 (s, 10H), 3.25 (s, 11H), 2.19 (s, 7H), 2.04 (s, 4H), 1.90 (s, 4H), 1.67 – 1.58 (m, 41H), 1.53 (s, 154H), 1.44 (m, 121H), 1.28 (m, 406H), 0.88 (t, *J* = 7.0 Hz, 55H).

**Deprotection of $\mathbf{Daba\text{-}CART_{Boc}}$ to form Daba-CART:**

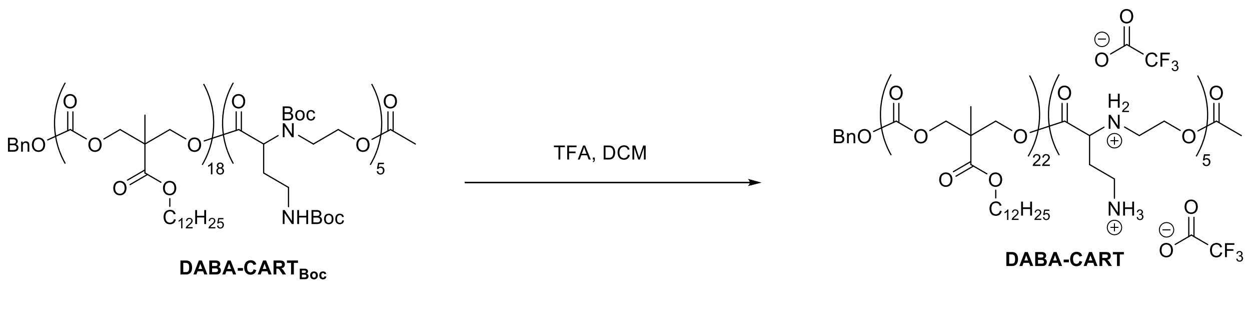


To a solution of $\mathbf{Daba\text{-}CART_{Boc}}$ (n = 18, m = 5, 32.4 mg) in dry, degassed dichloromethane (3.24 ml) was added trifluoroacetic acid (324 µl). The solution was stirred under a nitrogen atmosphere for 4 hours, after which point the product was concentrated to dryness to yield a clear oil (35.3 mg). The product **Daba-CART** was then dissolved in dimethyl sulfoxide to make a 5 mM solution, which was used in the in vivo experiments without further purification.

**Synthesis of Dapa-derived copolymer Dapa-CART$_{Boc}$**

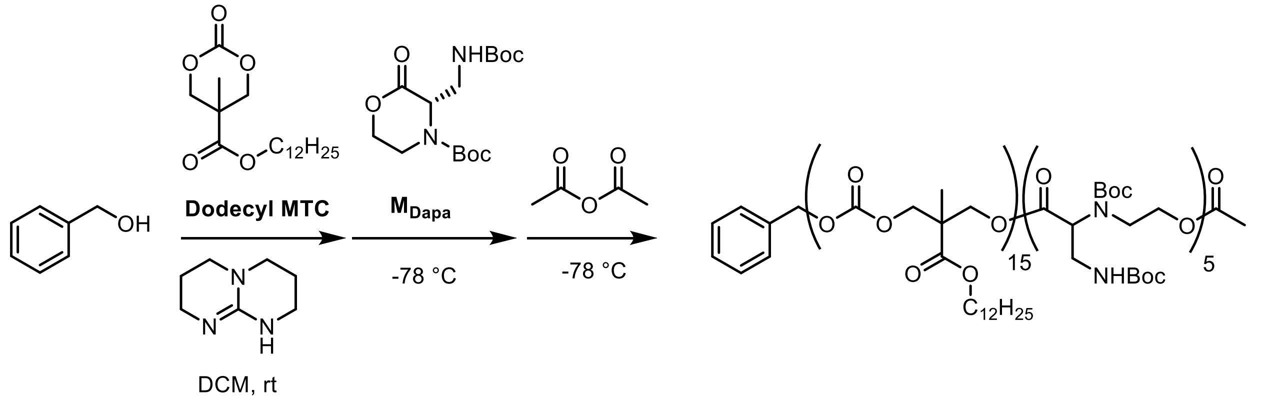


To a solution of **dodecyl MTC** (111.3 mg, 0.339 mmol, 16 eq) in dichloromethane (160 µl) was added a solution of benzyl alcohol (2.29 mg, 0.021 mmol, 1 eq) and 1,5,7-triazabicyclo[4.4.0]dec-5-ene (2.06 mg, 0.0014 mmol, 0.7 eq) in dichloromethane (20 µl) in a glovebox. This solution was stirred at room temperature. After 7 minutes, the solution of **dodecyl MTC**, initiator, and catalyst was added to predissolved **M$_{Dapa}$** (42.0 mg, 0.127 mmol, 6.0 eq) in dichloromethane (20uL). The solution was cooled to -78 $^{o}$C and stirred for 15 minutes, after which point a solution of acetic anhydride (0.5 mL, x.s.). This was allowed to warm to room temperature. The crude residue was redissolved in dichloromethane and then dialyzed (regenerated cellulose tubing, MWCO 1kD) against methanol for overnight to yield **dapa-CART$_{Boc}$** as a clear oil (109.4 mg). $^{1}$H NMR analysis (CDCl3, 600 MHz) reveals an oligomer with 15 dodecyl units and 5 dapa units. These DPs were determined by comparing the aromatic initiator signal (7.39-7.30 ppm, 5H) to the signal of the **M$_{Dapa}$** block (1.49-1.37 ppm, 88 H), and signal from dodecyl block (0.87 ppm, 44H). Note: in this manuscript, we define "**Dapa-CART**" as the oligomer with m = 15, n = 5; that is, BnO-D$_{15}$-*b*-Dapa$_{5}$-OAc.

GPC: $M_n$=3.9x10$^{3}$, Đ=1.28
1H NMR (600 MHz, cdcl3) δ 7.39 – 7.30 (m, 5H), 5.14 (s, 6H), 4.58 – 3.94 (m, 103H), 3.80 – 3.27 (m, 20H), 1.67 – 1.55 (m, 30H), 1.49 – 1.37 (m, 88H), 1.36 – 1.15 (m, 317H), 0.87 (t, J = 7.0 Hz, 44H).

**Deprotection of Dapa-CART$_{Boc}$**

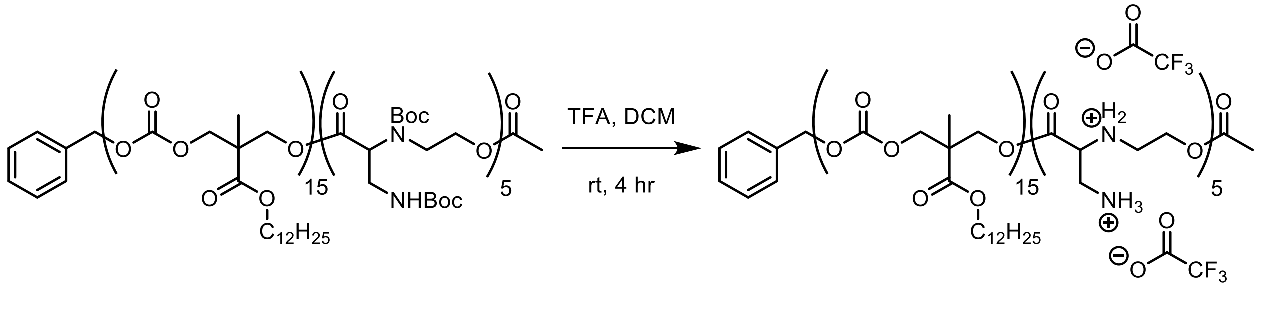


To a solution of Dapa-CART$_{Boc}$ ( 82 mg, 0.0122 mmol) in dry dichloromethane (5 mL) in a 2 dram scintilating vial charged with a stir bar was added trifluoroacetic acid (0.5 mL). The headspace was purged with nitrogen and the vial was capped. The reaction was stirred at room temperature for 4 hours. Upon completion the product was dried under reduced pressure affording a thin film (63.7 mg, 0.0093 mmol, 76% yield). The product was disolved in dimethylsulfoxide to make a 5 mM solution, which was filtered through a 200 nm PTFE syringe filter and stored at -20 celsius prior to in vivo experiments.

[1]H NMR (600 MHz, cdcl$_3$) δ 7.47 – 7.31 (m, 5H), 5.17 – 5.07 (m, 2H), 4.83 – 4.39 (m, 22H), 4.39 – 4.02 (m, 79H), 4.02 – 3.43 (m, 16H), 1.70 – 1.52 (m, 32H), 1.44 – 1.04 (m, 306H), 0.87 (t, *J* = 7.0 Hz, 43H).

**Table S1.** Physicochemical characterization of dicationic CARTs at +/- ratio of 10:1.

| CART | Z-average diameter (nm) | PDI | ζ potential (mV) | Encapsulation Efficiency (EE) % |
|---|---|---|---|---|
| D-Lys-CART | 210 ± 16 | 0.23 | +52 ± 5 | 95.74 ± 0.16 |
| D-Orn-CART | 183 ± 6 | 0.18 | +73 ± 1 | 96.00 ± 0.06 |
| D-Daba-CART | 170 ± 6 | 0.17 | +73 ± 1 | 95.84 ± 0.06 |
| D-Dapa CART | 180 ± 10 | 0.19 | +82 ± 4 | 96.23 ± 0.13 |

A

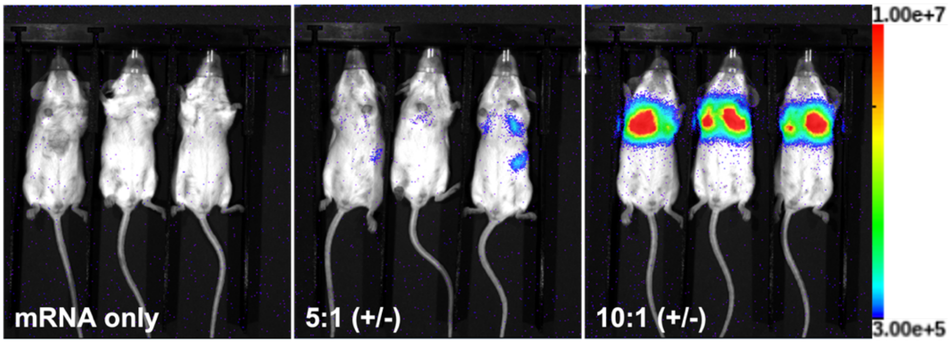

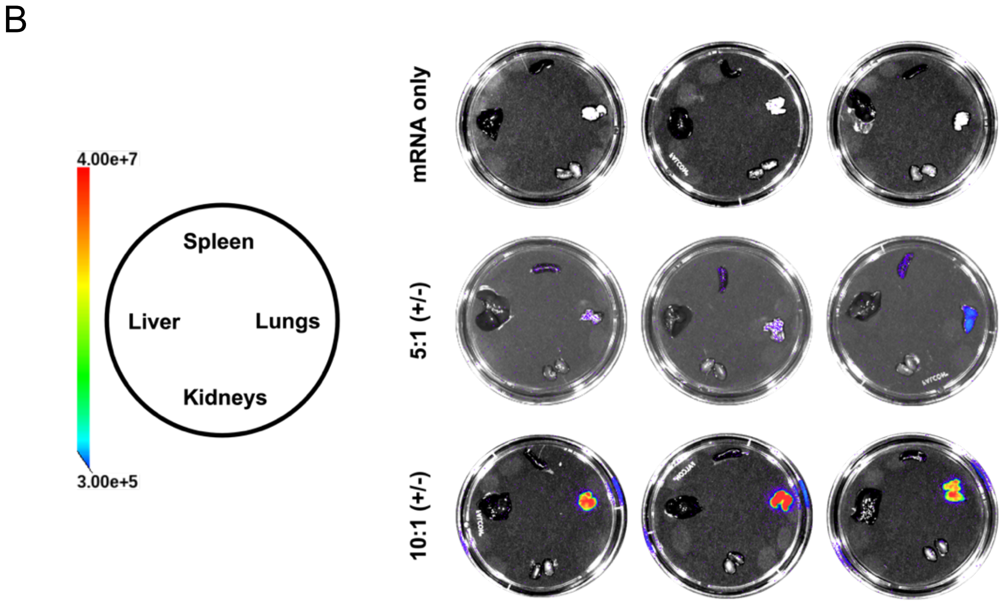


**Figure S2. Protein expression in organs of mice treated with Orn-CART fLuc mRNA particles.** Mice were injected intravenously with fLuc-encoding mRNA or with **Orn-CART** formulated with fLuc-encoding mRNA at 10:1 or 5:1 (+/-) charge ratios. (**A**) Full body and (**B**) organ bioluminescence images are shown for each condition.

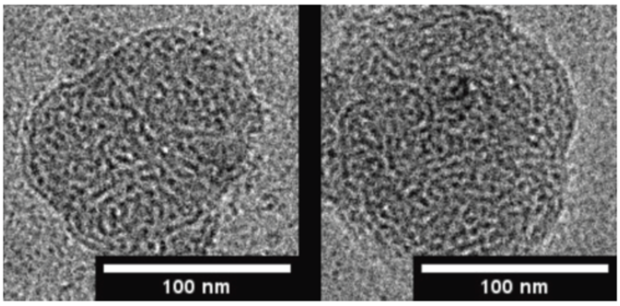


Orn-CART 10:1 (+/-) Orn-CART 5:1 (+/-)

**Figure S3.** Physicochemical characterization of $D_{15}$-$Orn_5$-CART/mRNA nanoparticles showing representative CryoEM micrographs of D-Orn-CART/mRNA nanoparticles with (+/-) charge ratios of 10:1 and 5:1 showing spherical particles with a worm-like internal morphology.

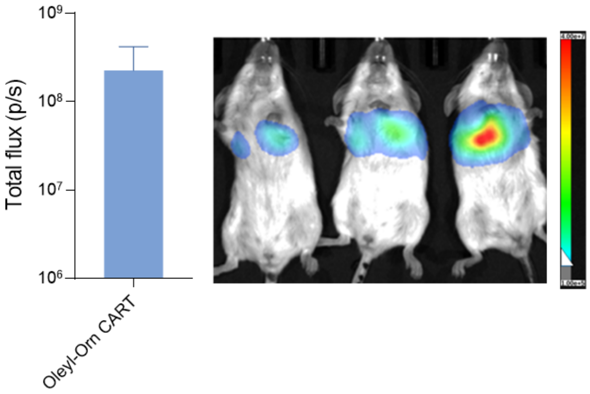


**Figure S4.** *In vivo* Bioluminesence of Oleyl-Orn CART. Mice were injected intravenously with 5µg fLuc-encoding mRNA or with Oleyl-**Orn-CART** formulated with fLuc-encoding mRNA at 10:1, *n* = 3 Independent biological replicates.

**Table S2.** Relative rates of Degradation of p(HE-AA) homopolymers (AA = Glycine, Lysine, Ornithine) in pH 6.5 phosphate-buffered water. Degradation rates reported as half life $t_{1/2}$

| | Glycine p(HE-Gly) | Lysine p(HE-Lys) | Ornithine p(HE-Orn) |
|---|---|---|---|
| $t_{1/2}$ | 3.5 min | 12 min | 16 min |

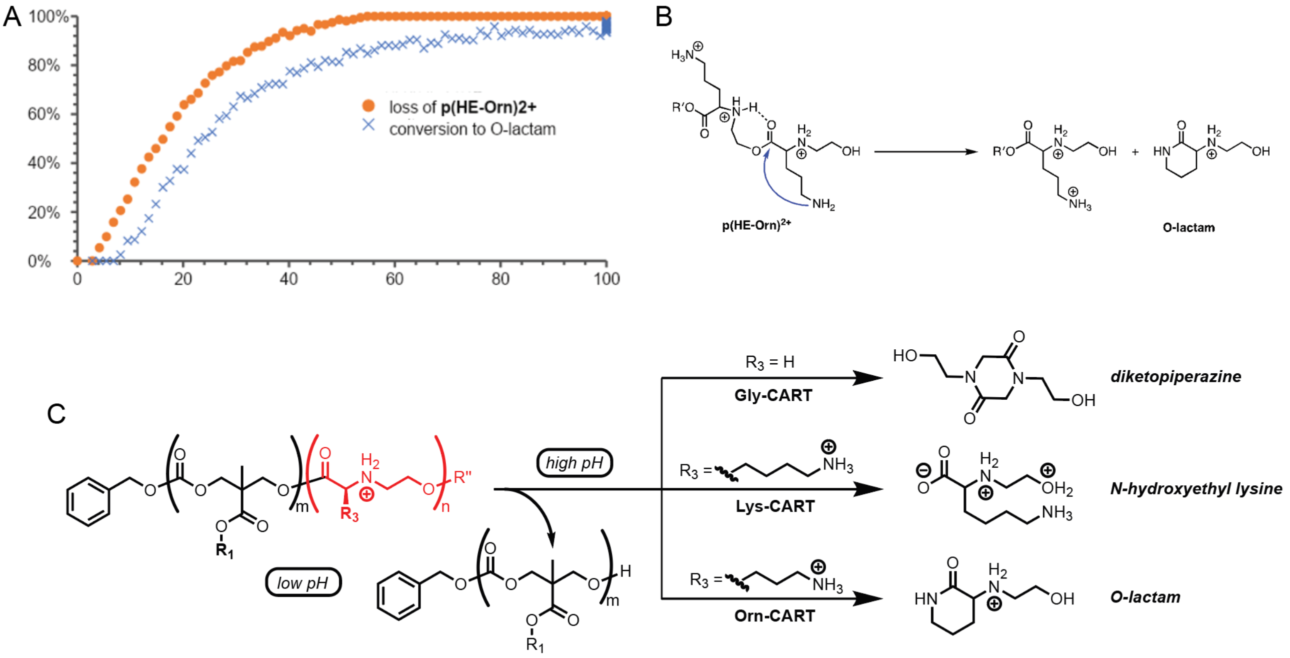


**Figure S5.** (A) Kinetics of rearrangement of $p(HE\text{-}Orn)^{2+}$ at pH 6.5. $p(HE\text{-}Orn)^{2+}$ (DP=21) was dissolved in $D_2O$ before the addition of phosphate-buffered $D_2O$ (pH6.5). (A) $^1H$ NMR spectra acquired at various timepoints revealed a $t_{1/2}$ of 12 min, with O-lactam to being major product and $(HE\text{-}Orn)^{2+}$ a negligible product. (B) Proposed degradation mechanism of Orn-CARTs. (C) Comparison between degradation mechanisms of D-Gly, D-Lys and D-Orn CARTs.

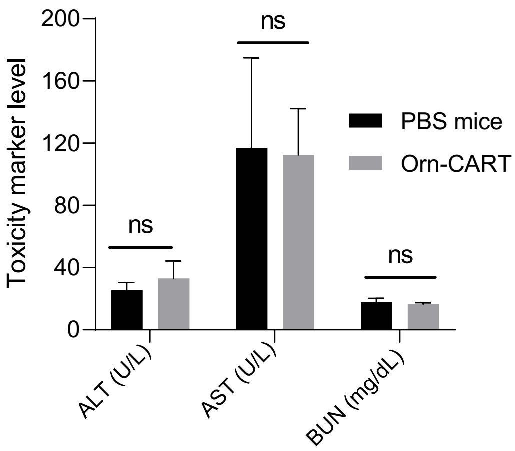


**Figure S6**. Blood chemistry analysis (ALT, AST, and BUN) of blood drawn from mice treated with $D_{15}Orn_5$-CART/mRNA NPs following intraveneous administration of 5µg Fluc mRNA. Data presented as mean ± SD, (ns; not-significant, *n = 3*).

**Table (S3)**. **Chronic toxicity study of D-Orn-CART**. Mice were injected on day 0, 7, 14 and 21 with 5µg Fluc mRNA Blood was collected 24 hr after the last dose, n = 3

| Marker | PBS control | D-Orn-CART | Normal range | Remarks |
|---|---|---|---|---|
| AST (mg/dL) | *616±377 | 281±99 | 184-220 | *Hemolysis observed |
| ALT (U/L) | 73±29 | 39±5 | 192-388 | |
| ALP (IU/L) | 140±4 | 132±13 | 171-183 | |
| BUN (mg/dL) | 22±2 | 23±1 | 20.3-24.7 | |

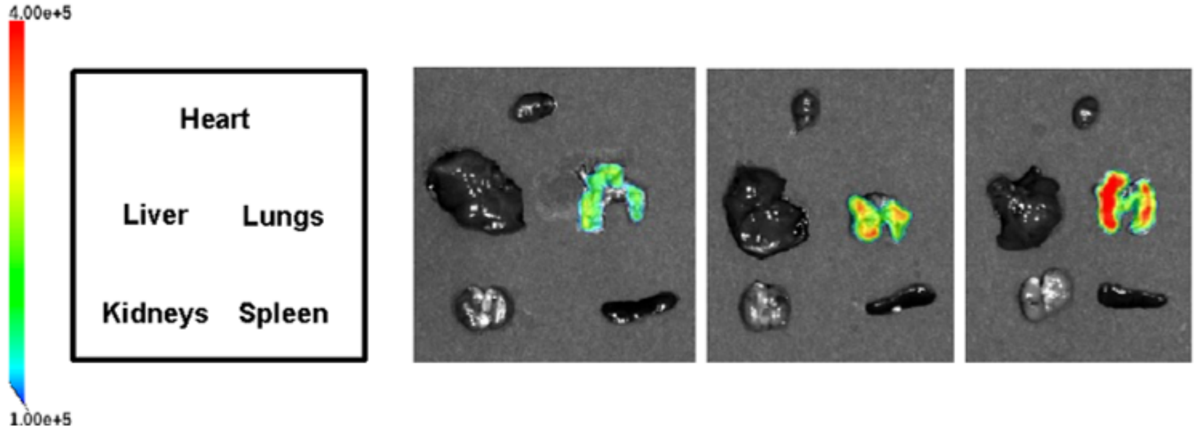


**Figure S7.** Protein expression in organs of mice removed 94 hours after treatment. Organ bioluminescence images.

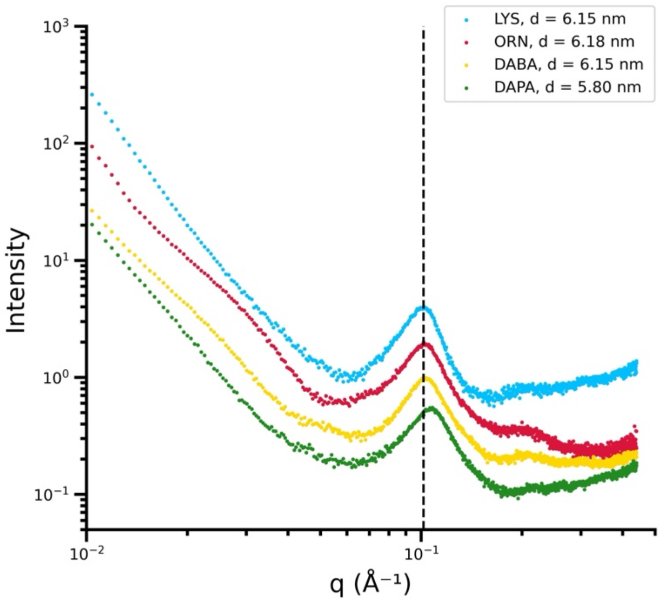


**Figure 8.** Small-Angle X-ray scattering profiles of CART/fLuc mRNA nanoparticles formulated at 10:1 +/- ratios of the dicationic CARTs. CARTs derived from D-Lys (blue), D-Orn (red), and D-Daba (yellow) have a peak at q ≈ 0.10 Å-1 which corresponds to a feature of 6.2 nm. D-Dapa (green) has a feature at at q ≈ 0.11 Å-1 corresponding to a feature around 5.8 nm.

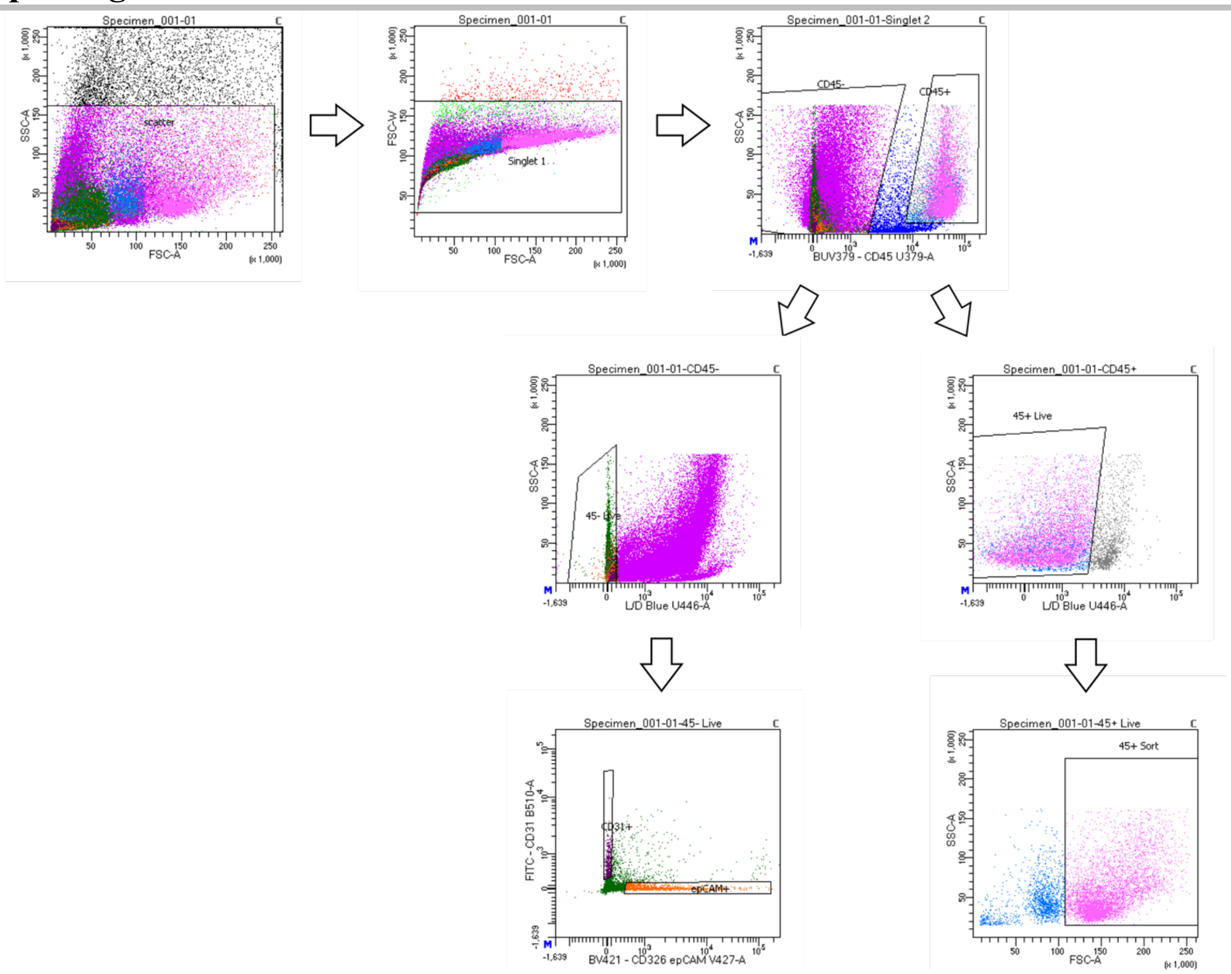


**Figure S9.** FACS gating strategy used for sorting lung epithelial, endothelial, and immune cell populations.

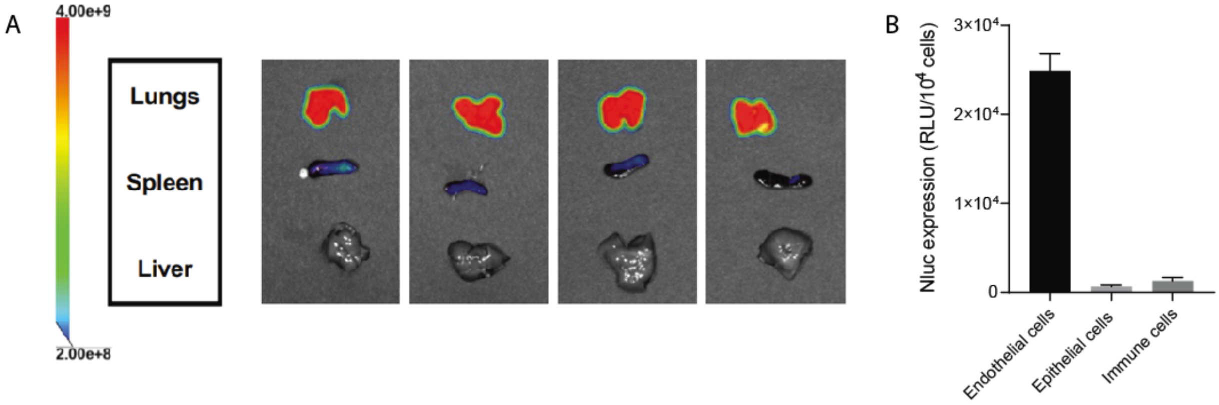


**Figure S10.** A) Organ bioluminescence images of mice treated with D-Orn-CART/NanoLuc mRNA nanoparticles. Mice were injected intravenously with D-Orn-CART formulated with NanoLuc-encoding mRNA at total dose of 10μg and a 10:1 (+/-) charge ratio, $n$ = 4 mice. B) Luminescence of lung cell subpopulations from mice treated intravenously with 10μg NanoLuc-encoding mRNA formulated with D-Orn-CART at a 10:1 (+/-) charge ratio, $n$ = 3.

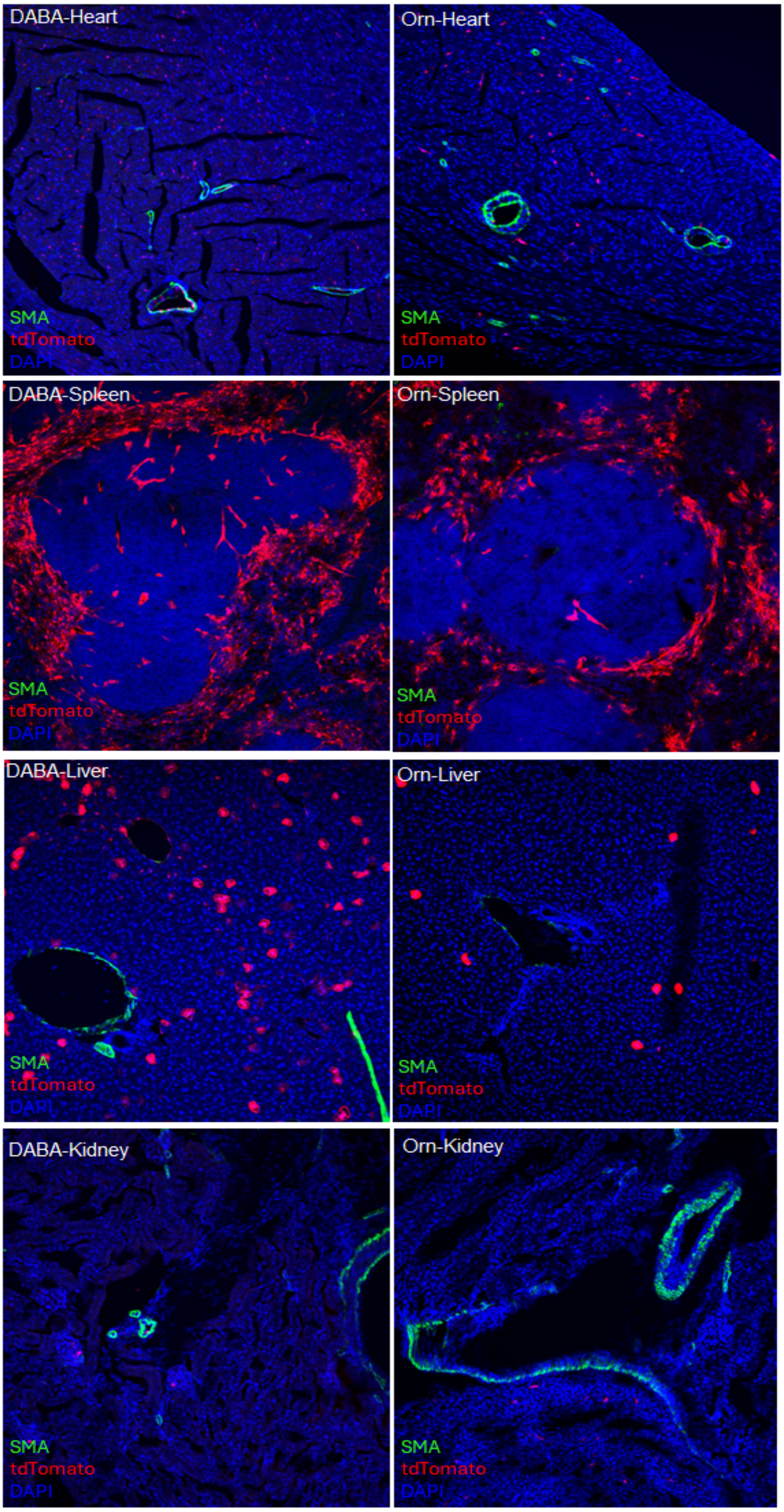


**Figure S11**. CART is localized to heart, spleen, liver and kidneys. Immunohistochemistry demonstrating positive tdTomato signal, indicating successful recombination in these organs from both DABA-CART and Orn-CART injections. SMA indicates smooth muscle α-actin. DAPI indicates 4′,6-diamidino-2-phenylindole.

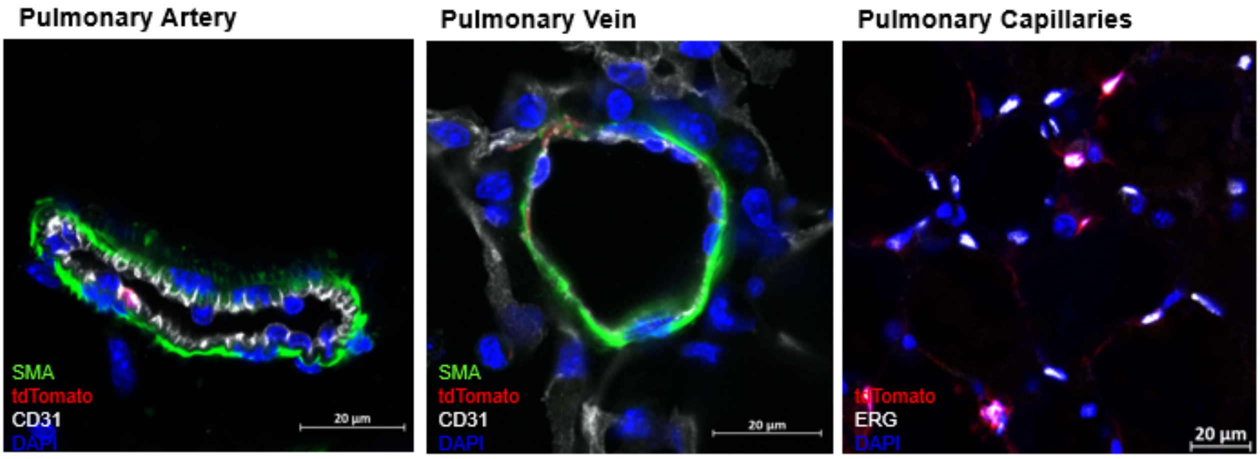


**Figure S12.** Lys-CART targets endothelial cells in the pulmonary vasculature showing minimal labelling of pulmonary arteries, pulmonary veins and pulmonary capillaries.

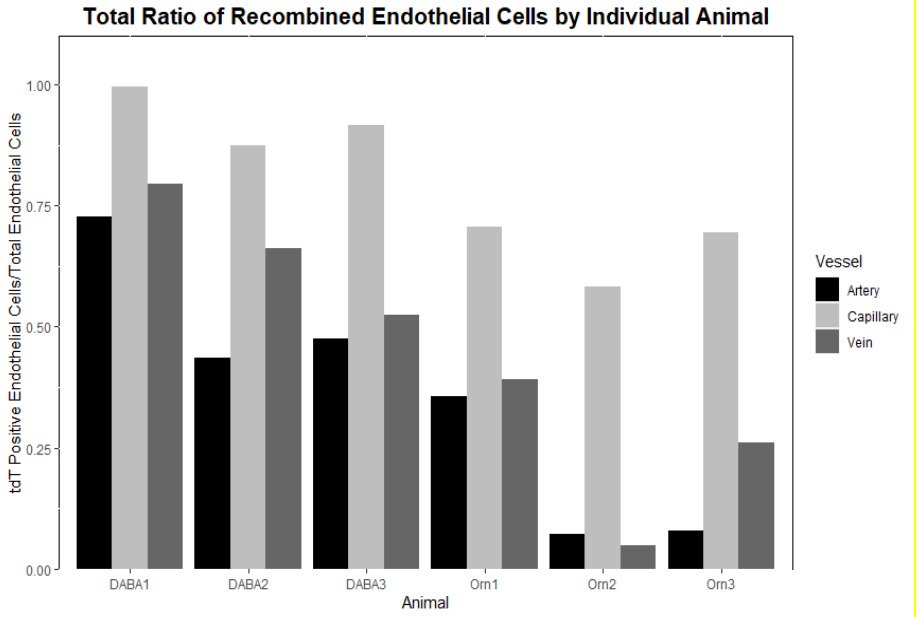


**Figure S13.** Variation in CART recombination efficiency between animals in DABA-CART ($p<0.01$) and Orn-CART ($p<0.01$) treated animals.

#### NMR spectra

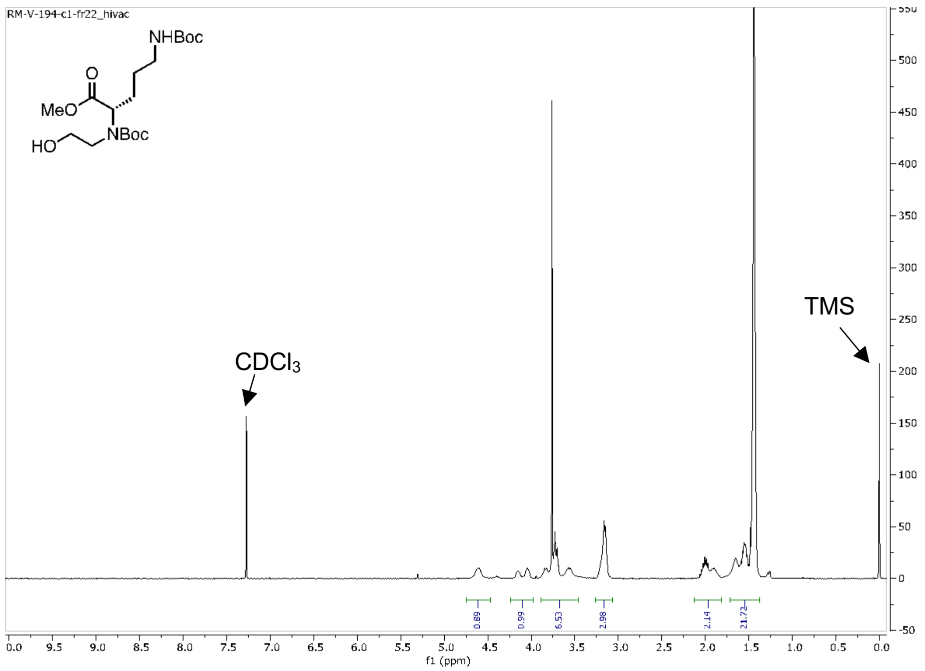


**Figure S14**. $^{1}H$ NMR spectrum of methyl 2-((*tert*-butoxycarbonyl)(2-hydroxyethyl)amino)-5-((*tert*-butoxycarbonyl)amino)pentanoate in $CDCl_3$.

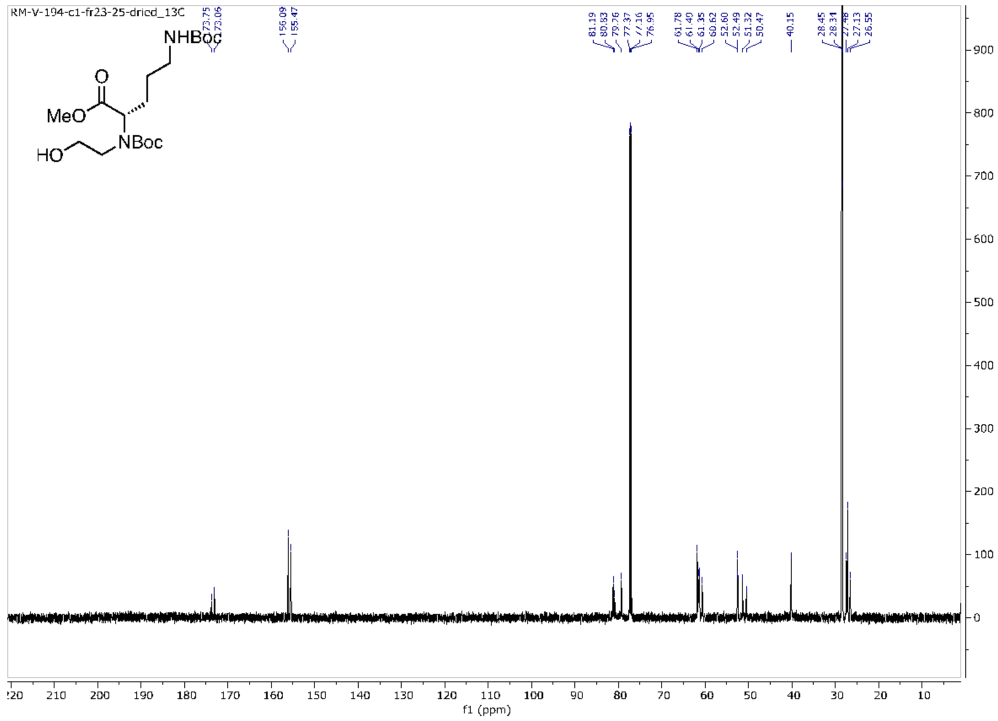


**Figure S15**. $^{13}C$ NMR spectrum of methyl 2-((*tert*-butoxycarbonyl)(2-hydroxyethyl)amino)-5-((*tert*-butoxycarbonyl)amino)pentanoate in $CDCl_3$.

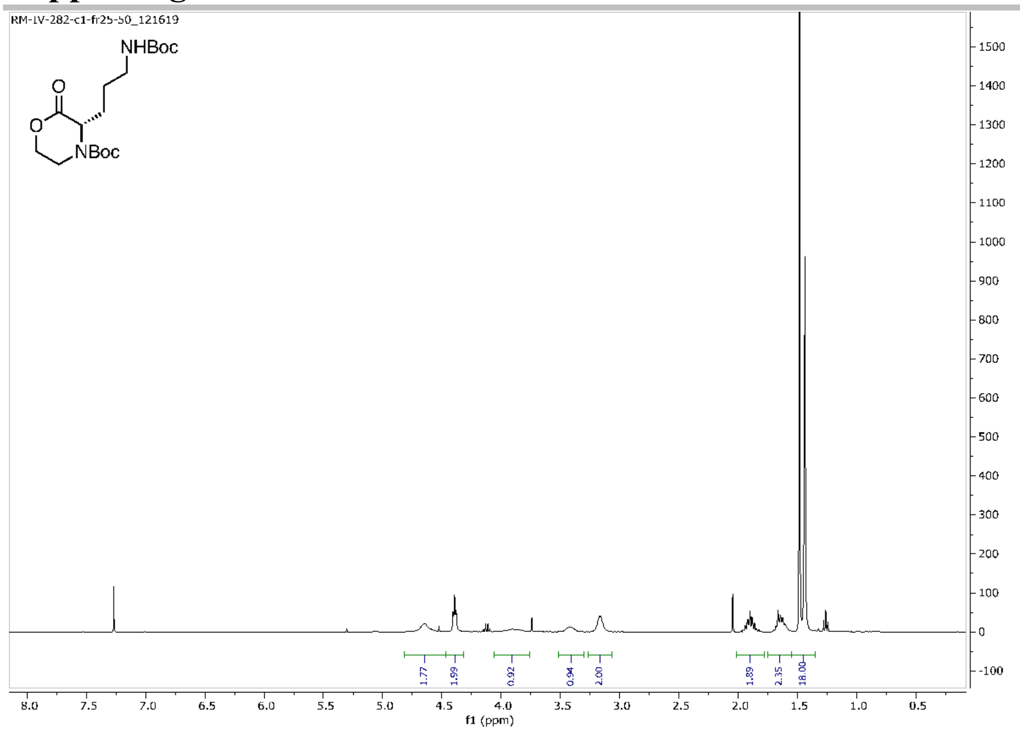


**Figure S16.** $^{1}H$ NMR spectrum of *tert*-butyl 3-(3-((*tert*-butoxycarbonyl)amino)propyl)-2-oxomorpholine-4-carboxylate ($M_{Orn}$) in $CDCl_3$.

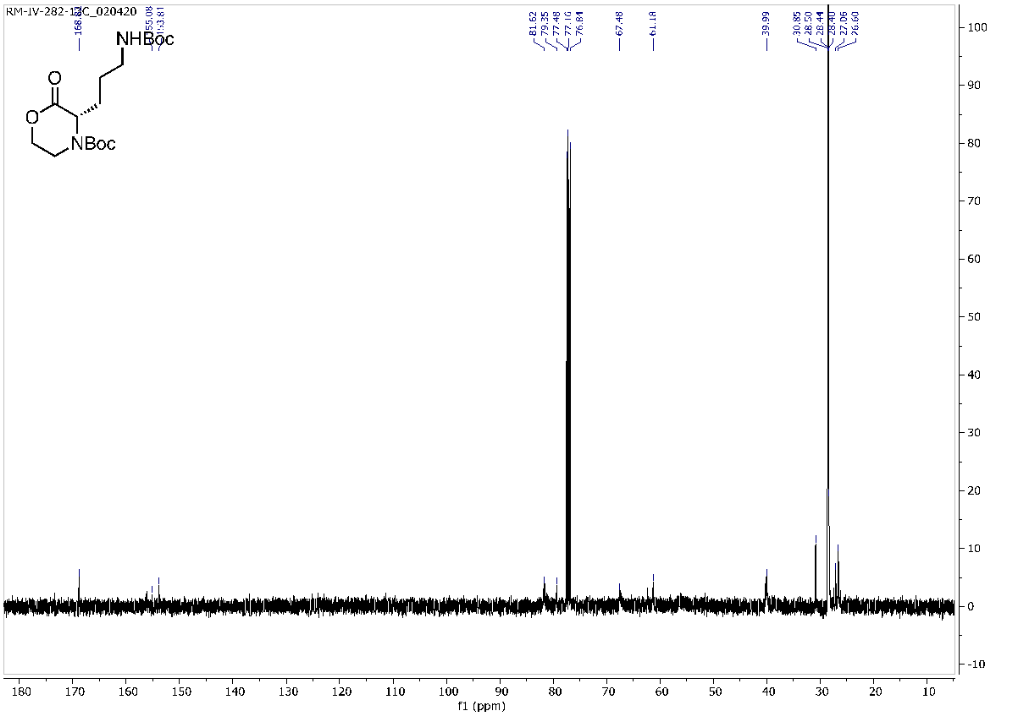


**Figure S17.** $^{13}C$ NMR spectrum of *tert*-butyl 3-(3-((*tert*-butoxycarbonyl)amino)propyl)-2-oxomorpholine-4-carboxylate ($M_{Orn}$) in $CDCl_3$.

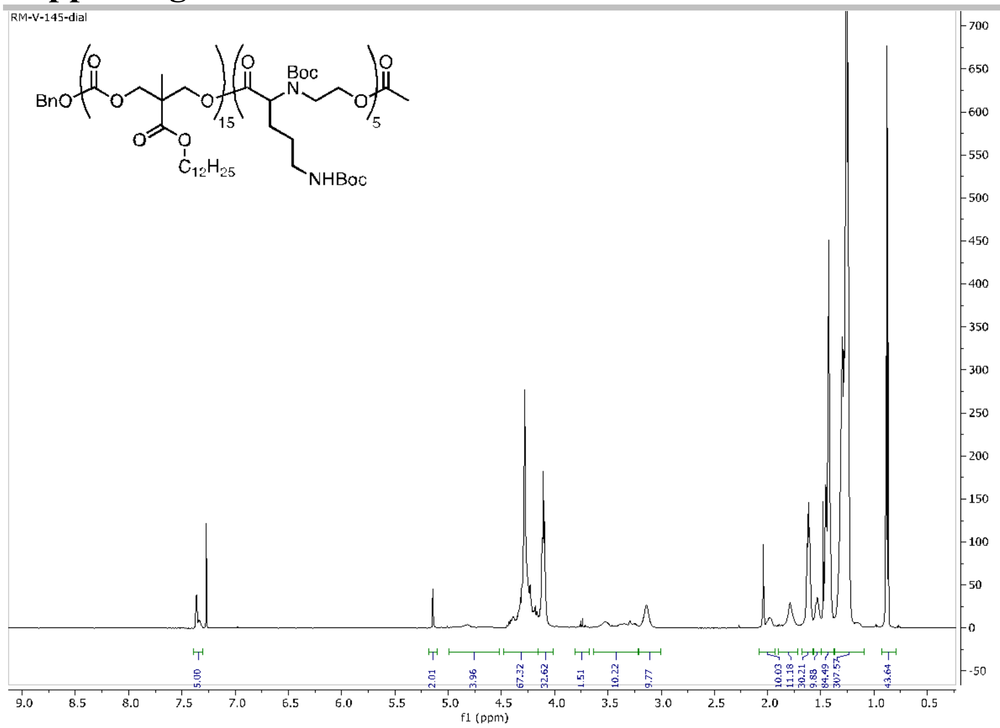

**Figure S18**. $^{1}H$ NMR spectrum of Orn-$CART_{Boc}$ in $CDCl_3$.

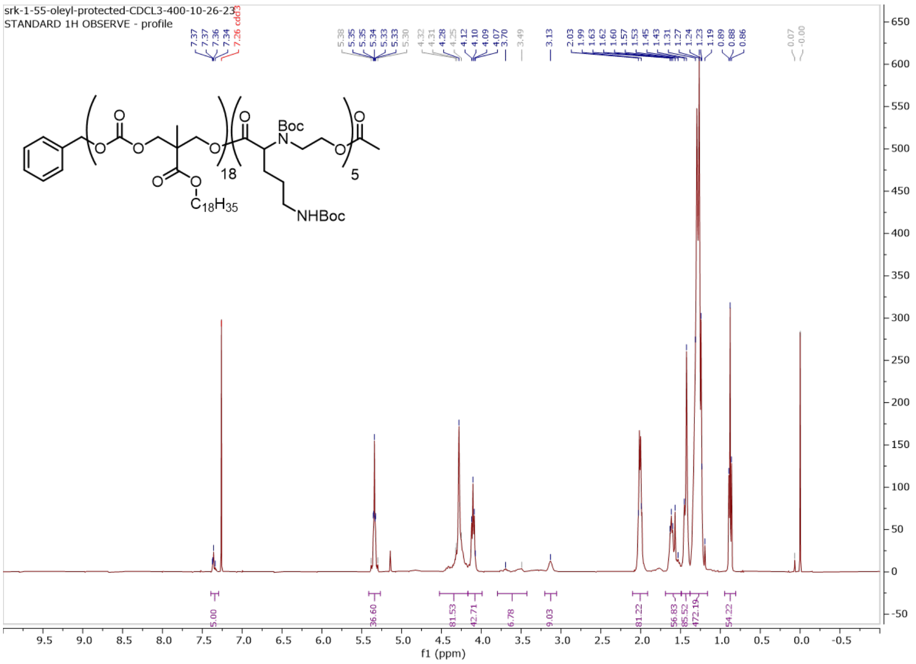

**Figure S19**. $^{1}H$ NMR spectrum of Oleyl Orn-$CART_{Boc}$ in $CDCl_3$.

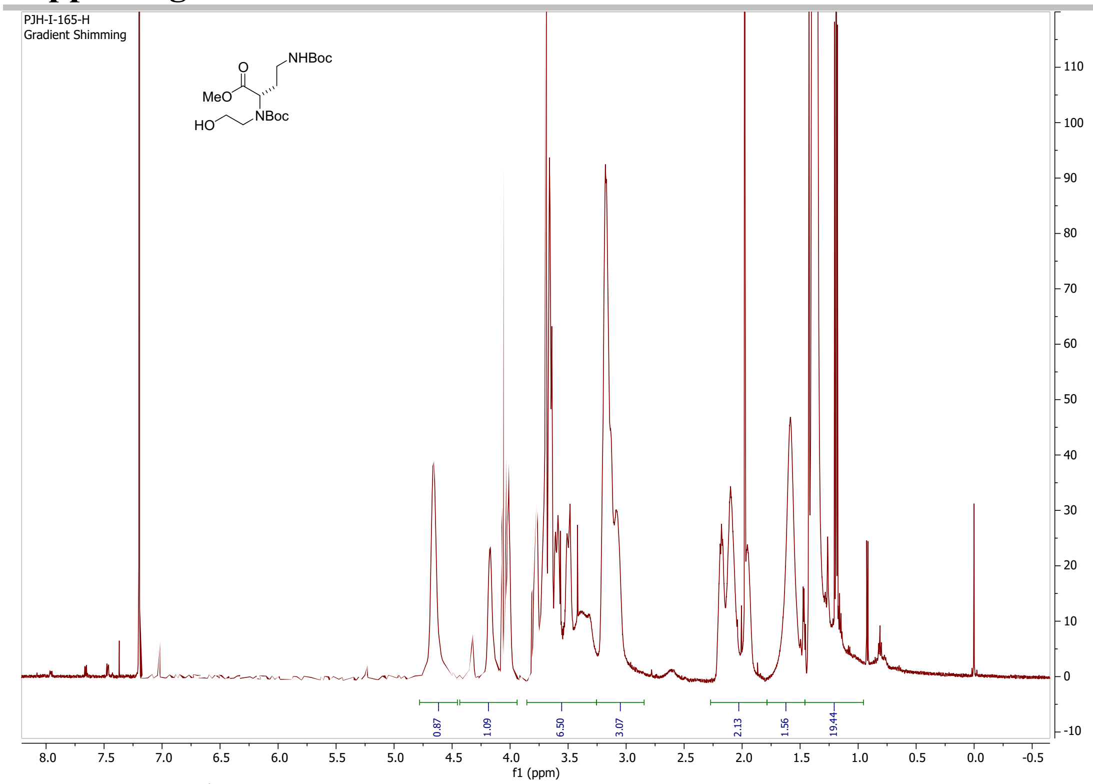

**Figure S20**. [1]H NMR spectrum of Daba monomer percursor in $CDCl_3$.

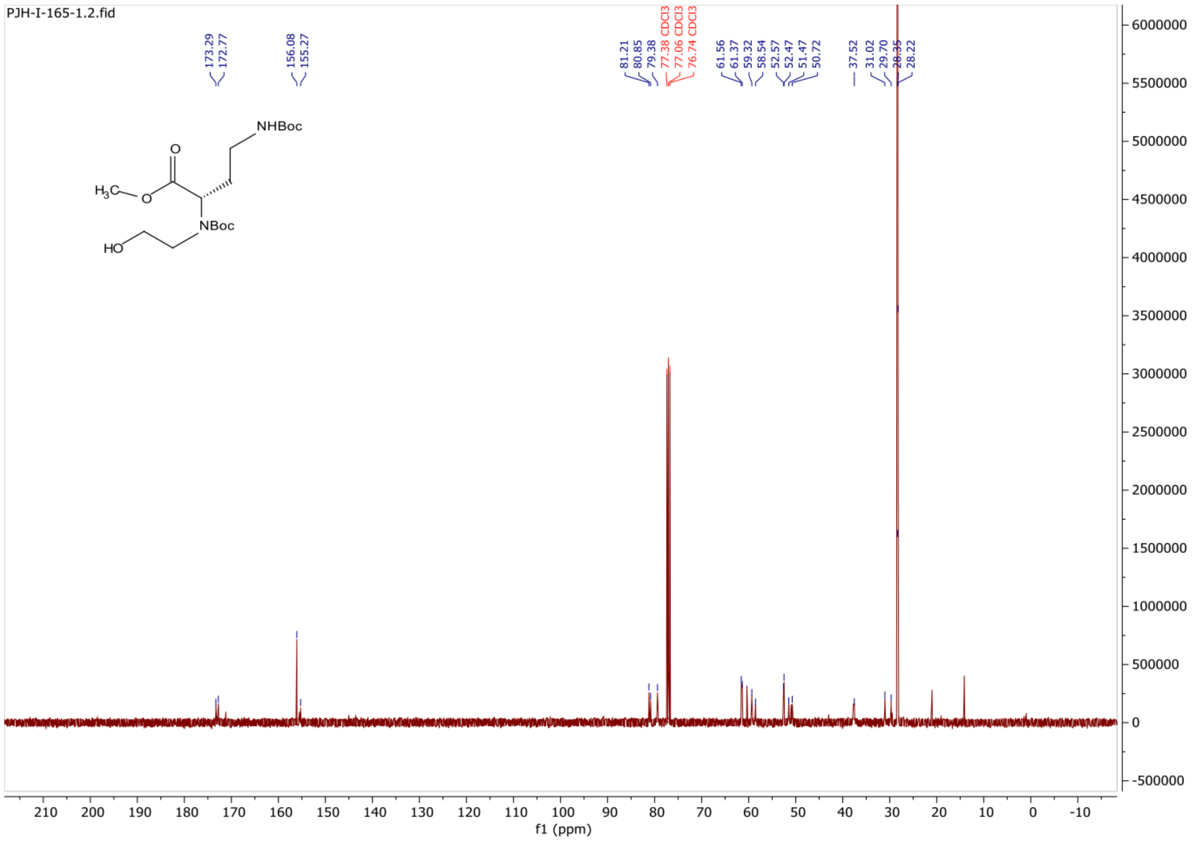

**Figure S21**. [13]C NMR spectrum of Daba monomer percursor in $CDCl_3$.

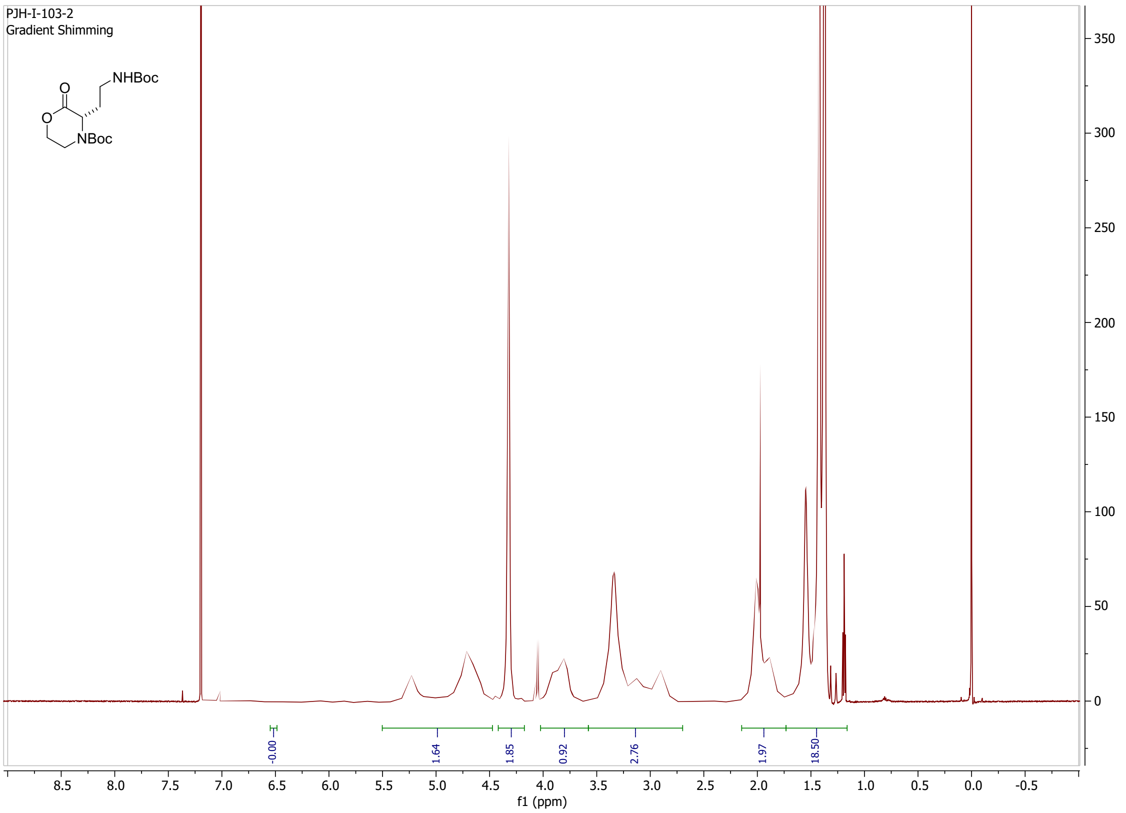


**Figure S22**. [1]H NMR spectrum of Daba monomer in $CDCl_3$.

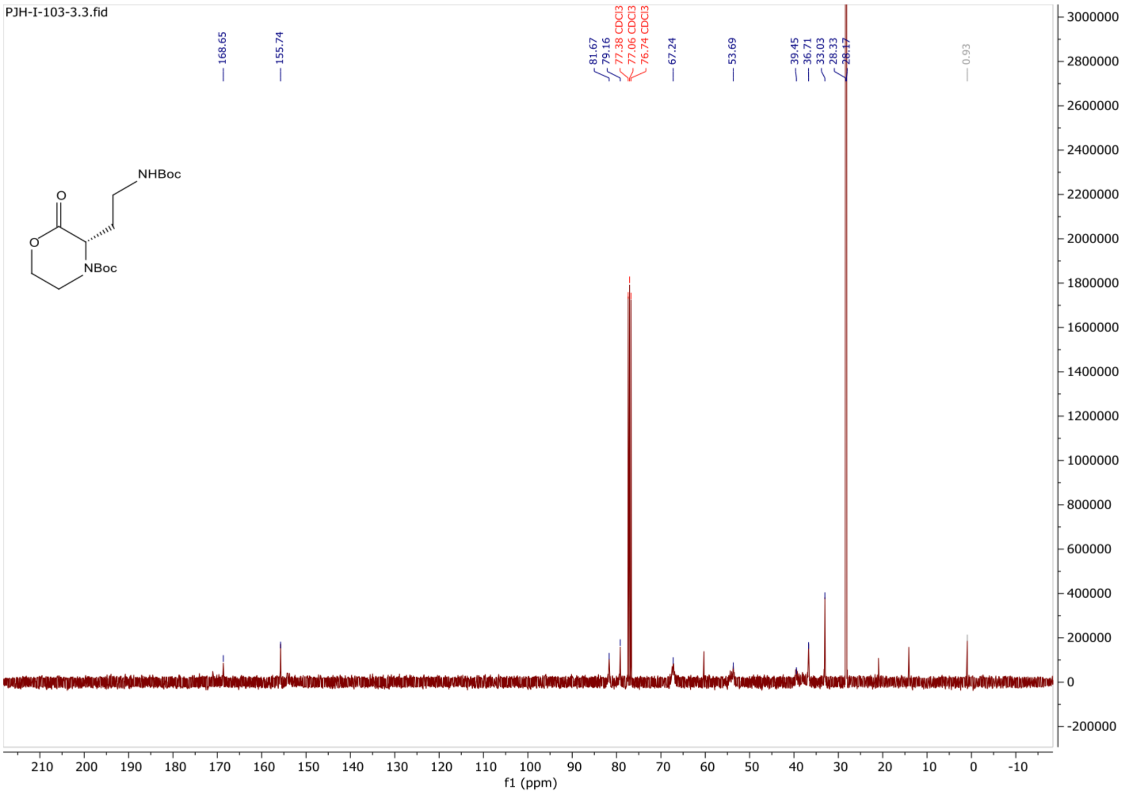


**Figure S23**. [13]C NMR spectrum of Daba monomer in $CDCl_3$.

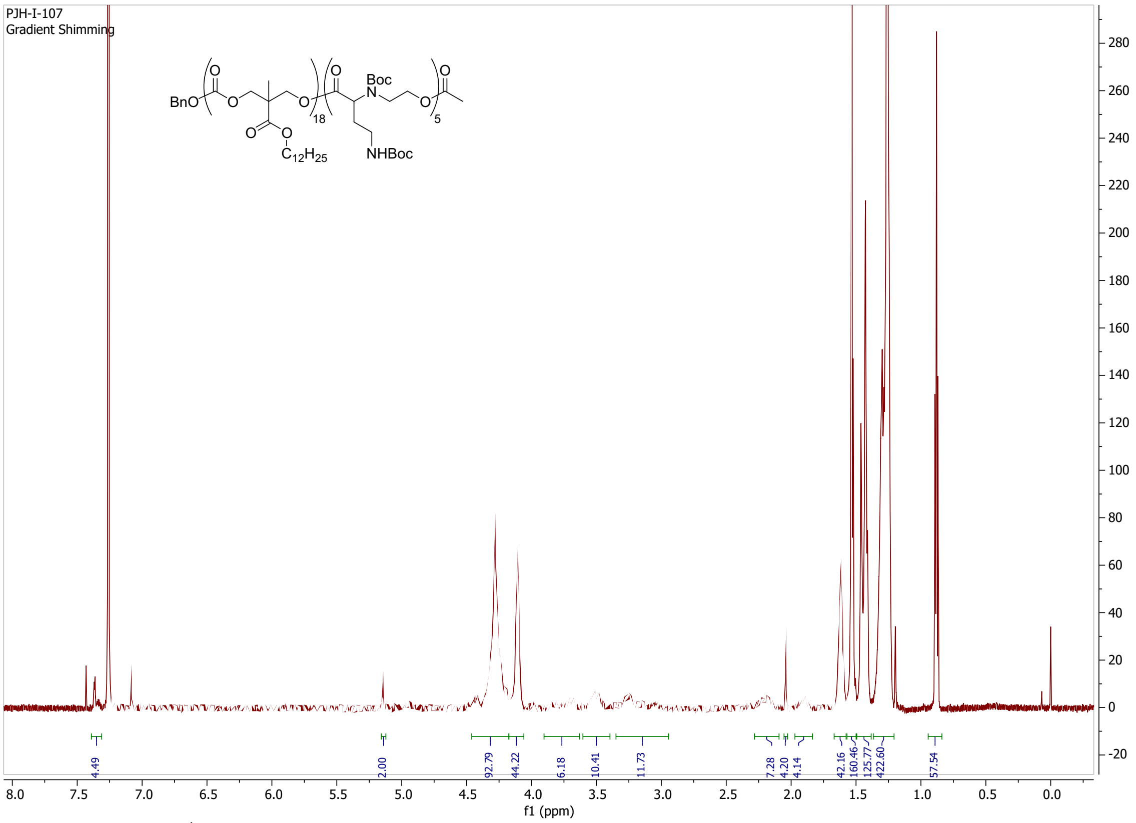


**Figure S24**. [1]H NMR spectrum of Daba-CART$_{Boc}$ in $CDCl_3$.

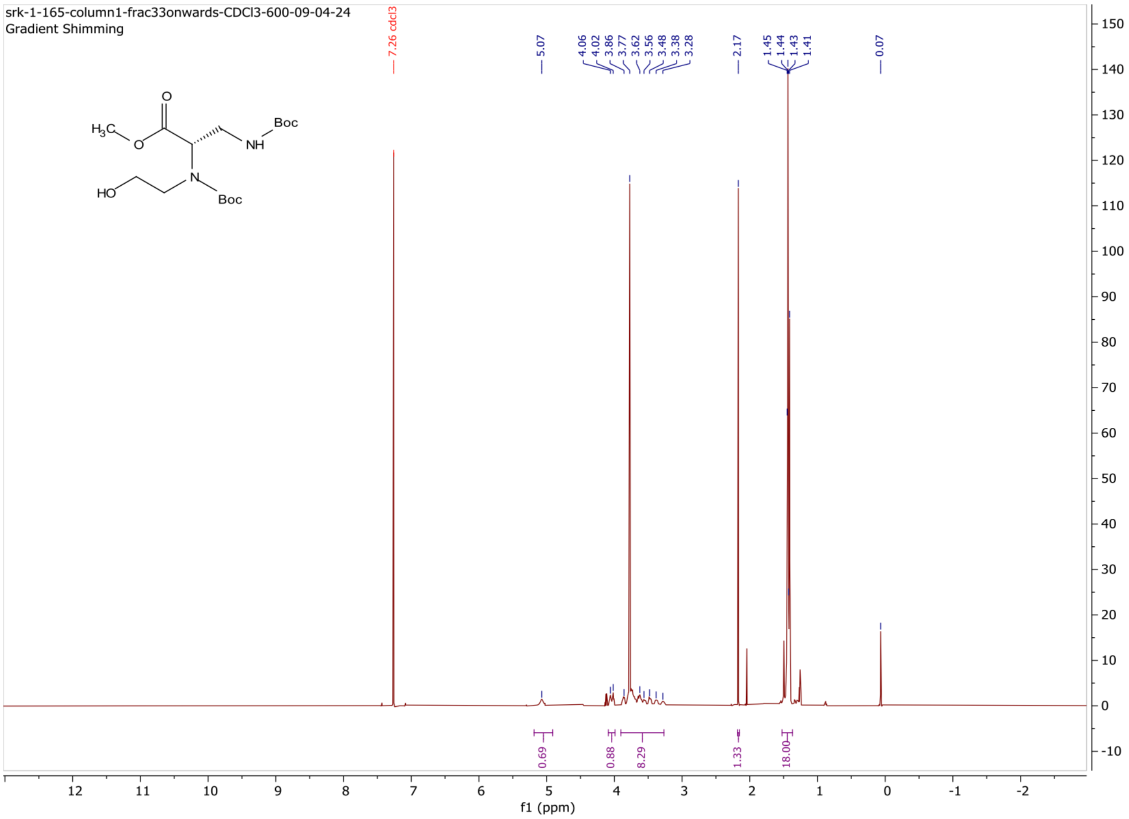


**Figure S25**. [1]H NMR spectrum of Dapa monomer precurser in $CDCl_3$.

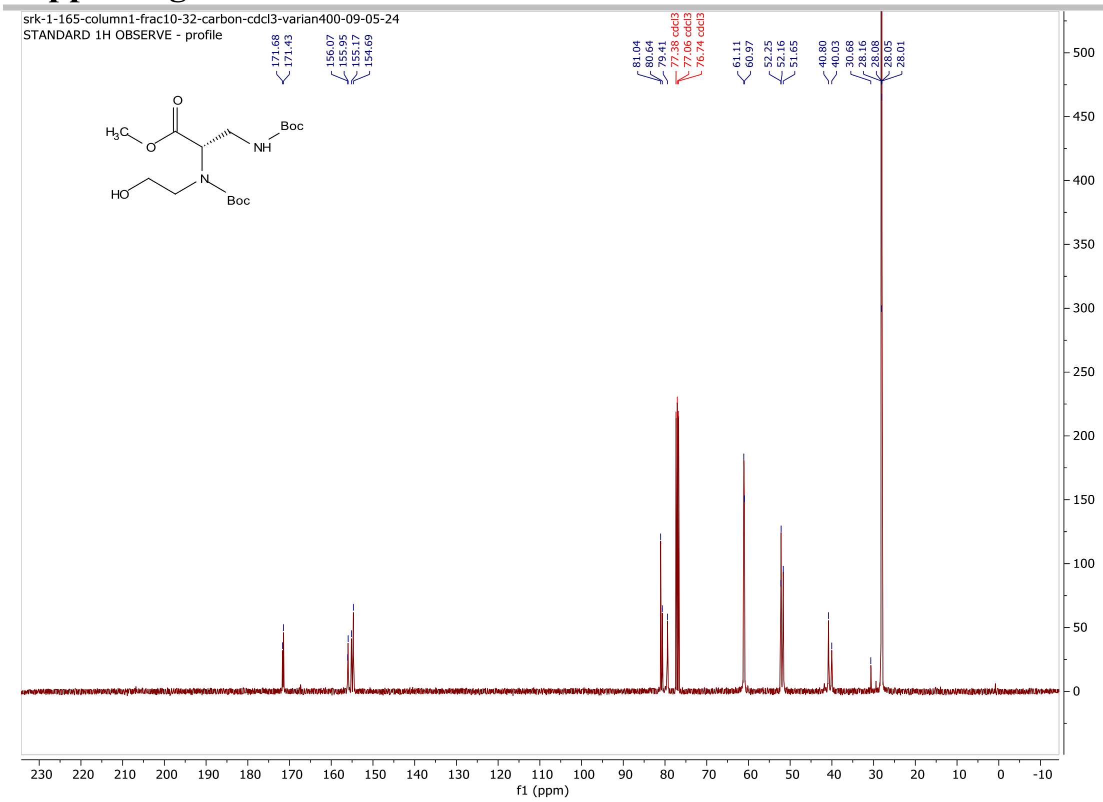


**Figure S26**. $^{13}$C NMR spectrum of Dapa monomer precurser in $CDCl_3$.

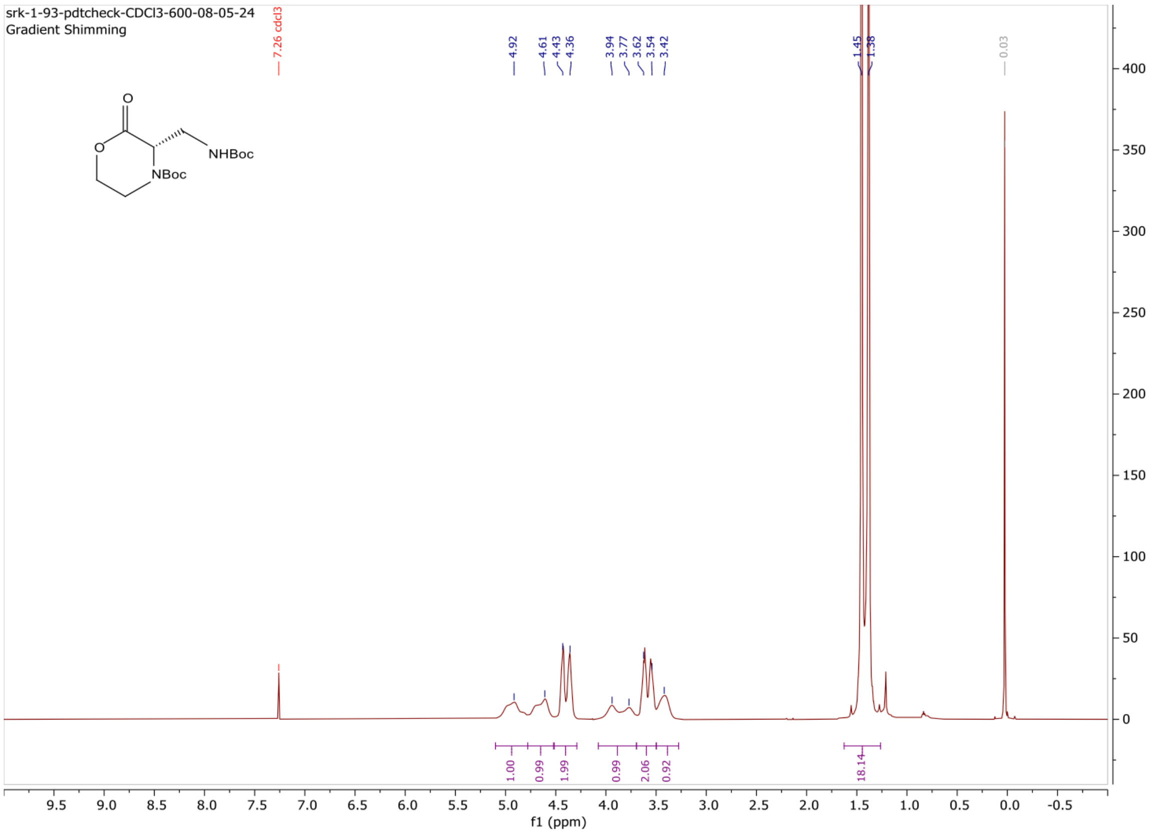


**Figure S27.** $^{1}H$ NMR spectrum of $M_{Dapa}$ in $CDCl_3$

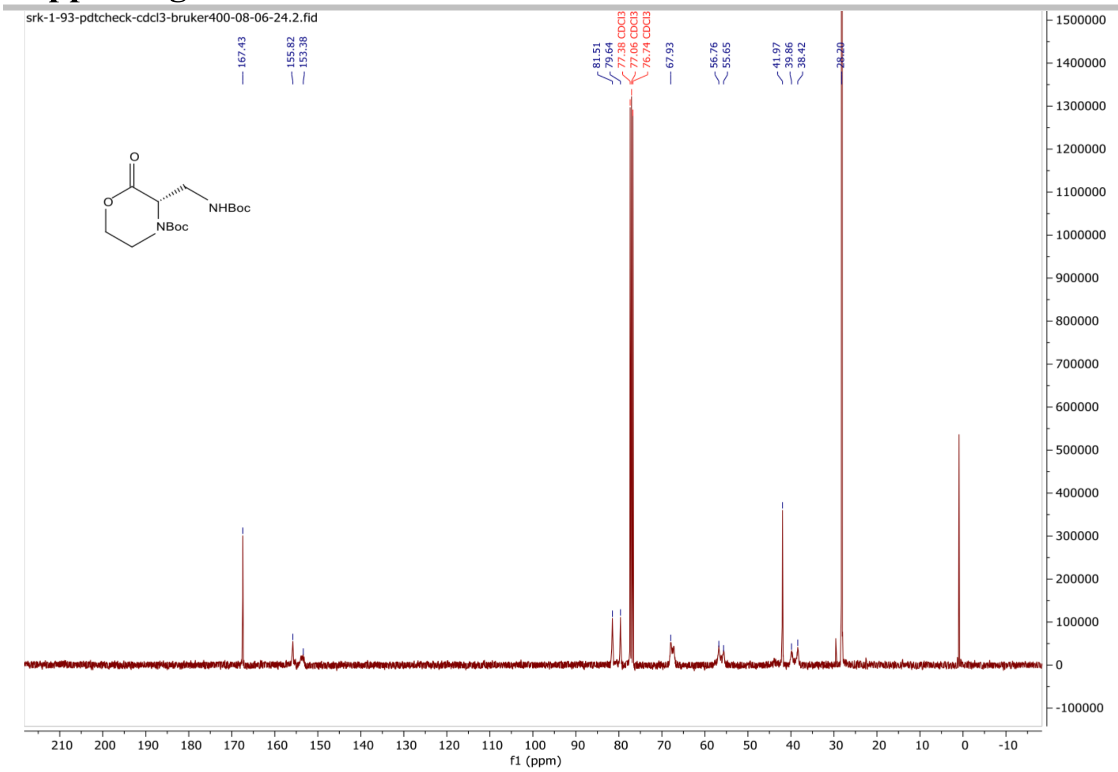


**Figure S28.** $^{13}C$ NMR spectrum of $M_{Dapa}$ in $CDCl_3$

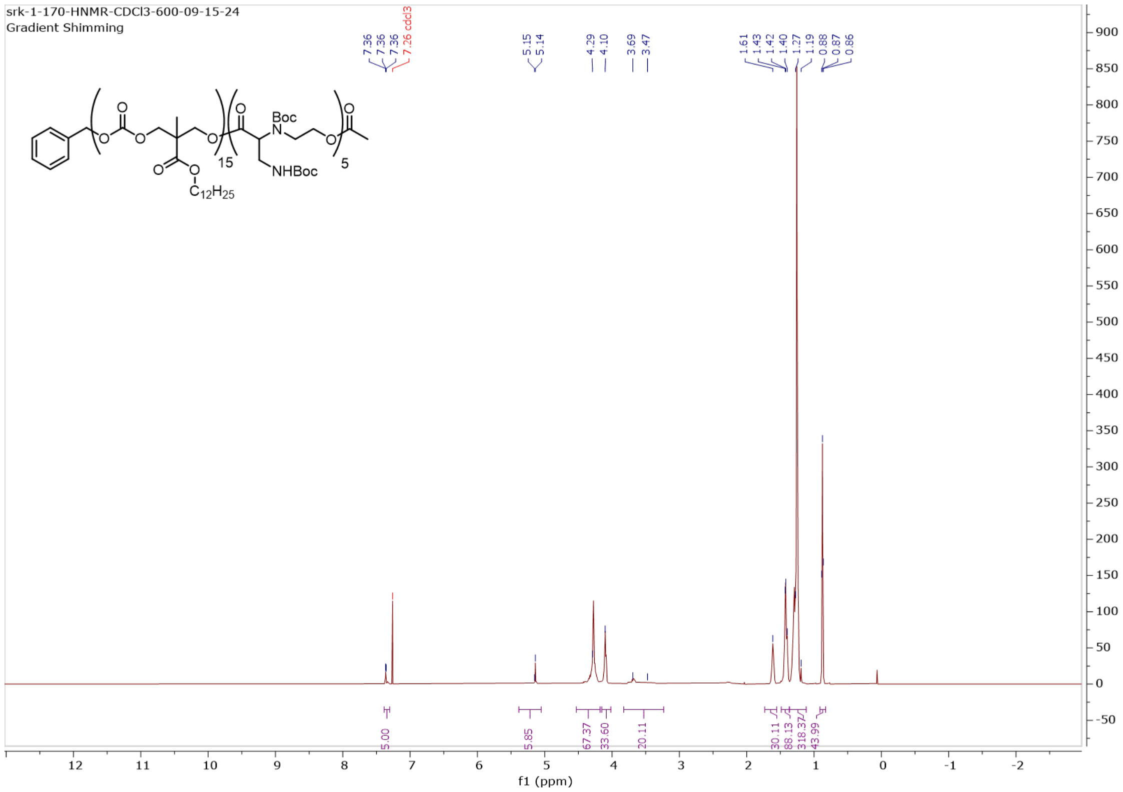


**Figure S29**. $^{1}H$ NMR spectrum of Dapa-$CART_{Boc}$ in $CDCl_3$.

#### References

1. Pratt, R. C.; Nederberg, F.; Waymouth, R. M.; Hedrick, J. L., Tagging alcohols with cyclic carbonate: a versatile equivalent of (meth)acrylate for ring-opening polymerization. *Chemical Communications* **2008**, (1), 114-116.

2. McKinlay, C. J.; Benner, N. L.; Haabeth, O. A.; Waymouth, R. M.; Wender, P. A., Enhanced mRNA delivery into lymphocytes enabled by lipid-varied libraries of charge-altering releasable transporters. *Proceedings of the National Academy of Sciences* **2018**, 201805358.

3. Kumar ME, Bogard PE, Espinoza FH, Menke DB, Kingsley DM, Krasnow MA. Mesenchymal cells. Defining a mesenchymal progenitor niche at single-cell resolution. *Science*. 2014;346:1258810. doi: 10.1126/science.1258810
